# *In vivo* CRISPRi screen reveals the differential requirement for mitochondrial respiratory chain function between *in vivo* and *in vitro* tumor growth

**DOI:** 10.1101/2021.10.03.462937

**Authors:** Hiroki Nakaoka, Neal Bennett, Ross A. Okimoto, Danny Laurent, Yoshitaka Sei, Trever Bivona, Johanna ten Hoeve, Thomas G. Graeber, Ken Nakamura, Jean L. Nakamura

**Author notes:** Address correspondence to: Jean Nakamura, University of California, San Francisco, Helen Diller Family Cancer Research Building, 1450 Third Street, San Francisco, CA 94158. These authors contributed equally (Hiroki Nakaoka, Neal Bennett).

## Abstract

The Warburg effect, aerobic glycolysis, is a hallmark feature of cancer cells grown in culture. However, the relative roles of glycolysis and respiratory metabolism in supporting *in vivo* tumor growth and significant processes such as tumor dissemination and metastases remain poorly understood, particularly on a systems level. Using a CRISPRi mini-library enriched for mitochondrial ribosomal protein and respiratory chain genes in multiple human lung cancer cell lines we analyzed *in vivo* metabolic requirements in xenograft tumors grown in distinct anatomic contexts. While knockdown of mitochondrial ribosomal protein and respiratory chain genes (mito-respiratory genes) has little impact on growth *in vitro*, tumor cells depend heavily on these genes when grown *in vivo* as either flank or primary orthotopic lung tumor xenografts. In contrast, respiratory function is comparatively dispensable for metastatic tumor growth. RNA-Seq and metabolomics analysis of tumor cells expressing individual sgRNAs against mito-respiratory genes indicate overexpression of glycolytic genes and increased sensitivity of glycolytic inhibition compared to control when grown *in vitro*, but when grown *in vivo* as primary tumors these cells downregulate glycolytic mechanisms. These studies demonstrate that discrete perturbations of mitochondrial metabolism impact *in vivo* tumor growth in a context-specific manner and provides systems-level evidence that respiratory function modulates tumor growth *in vivo*, suggesting that ATP limits growth and metastatic potential.

## INTRODUCTION

The dysregulation of cellular energy metabolism is an early fundamental event in tumorigenesis and a hallmark of cancer. Cancer cells modulate their metabolism as they proliferate, outpace normal cells in growth, and establish disease in diverse and often nutrient-restricted environments. The Warburg effect observed in cancer cells refers to the preferential use of aerobic glycolysis, which produces less ATP than aerobic respiration while favoring biosynthetic functions necessary for tumor growth (1). The specific roles of ATP-modulating mechanisms in supporting tumor growth are poorly understood, particularly *in vivo*. Most studies investigating the Warburg effect are performed in cultured cells and the full role of metabolic processes *in vivo*, for example the requirement for mitochondrial-derived ATP, remains undefined, especially as it pertains to *in vivo* tumor growth. In contrast to *in vitro* tumor modeling, tumor growth *in vivo* tests specific physiologic contexts in which cancer cells must metabolically adapt to thrive. We sought to determine whether mechanisms that modulate cellular ATP also impact cancer cell growth *in vivo*, employing multiple models to test anatomic context-dependence.

In prior work, we developed a high throughput screening paradigm to identify genetic regulators of ATP, combining FACS and CRISPR with an ATP-FRET sensor capable of monitoring real-time changes in ATP concentrations within individual living cells (2). Using this sensor, we performed genome-wide CRISPRi and CRISPRa screens to define an “ATPome” of genes and pathways that regulate ATP levels through energy substrate-specific pathways (respiration or glycolysis)(3). A key finding is that many genes and pathways that preserve or reduce ATP exert these effects only *under specific metabolic conditions defined by substrate availability*. Glycolysis and respiration demonstrate cross-optimization on a systems level, that is, suppression of one metabolic mechanism (via members of specific gene classes) optimizes the alternative mechanism. Silencing genes required for respiratory-derived ATP modestly suppressed *in vitro* growth under respiratory conditions, but conversely increased *in vitro* tumor cell growth under glycolytic conditions. This evidence of cross-optimization also points to highly modulatable metabolic control measurable in cellular ATP; they also indicate a broader repertoire of mechanisms available to cells for optimizing their metabolic function.

Respiration and glycolysis-derived ATP production are influenced by gene expression within both metabolic mechanisms, but whether and how each of these mechanisms of ATP production differentially impact *in vivo* growth, especially when multiple anatomically separate tumor sites have developed, is not known. We developed a custom CRISPRi mini-library composed of sgRNAs against genes modulating cellular ATP levels (identified from the ATPome(3)), to test whether ATP levels correlate with *in vivo* growth of tumors in multiple pre-clinical mouse models. *In vivo* xenograft models revealed differential growth effects of ATP-modulating genes within discrete anatomic sites, where suppression of genes involved in mitochondrial-derived ATP associated with reduced tumor growth in primary tumor models. To evaluate the molecular basis for this reduced growth we performed RNA Seq and metabolomics analysis in isogenic lung cancer cells engineered with discrete silencing of mito-respiratory genes; these studies indicate profound differences in metabolism, particularly glycolytic metabolic profile, that is highly tumor growth context dependent.

To our knowledge, a systems-level analysis of genetic modulators of ATP combined with *in vivo* functional readouts has not been performed in human cancers. By evaluating tumor growth in multiple *in vivo* contexts, our findings illustrate critical differences in ATP requirements between *in vitro* and *in vivo* tumor growth, pointing to a functional role for ATP and mitochondrial processes that is specific to regional growth and distinguishes primary tumors from distant metastases.

## RESULTS

### Primary *In vivo* tumor growth requires mitochondrial-derived ATP

Lung cancer is the most common cause of death due to cancer in the United States and is characterized by primary solid tumors that can metastasize widely to diverse organs. HCC827 and H1975 human lung cancer cells are model cell lines for Epidermal Growth Factor Receptor (EGFR)-mutant human non-small cell lung cancer (4, 5). To determine whether ATP-modulating genes impact the *in vivo* tumor growth of lung cancers, we transduced HCC827 cells expressing dCas9-KRAB with a custom CRISPRi mini-library enriched with respiratory and glycolytic hits that most significantly influenced ATP levels (high or low, depending on the substrate conditions). This mini-library contains over 400 sgRNAs (1/5^th^ of which are non-targeting sgRNAs included as negative controls), with multiple sgRNAs against each gene target (typically two to four unique sgRNAs/gene, sgRNAs listed in Supplementary Table 1) (3).

We postulated that similar to *in vitro* growth, individual gene silencing *in vivo* could confer relative growth advantages or disadvantages that could be estimated on the basis of sgRNA representation. After antibiotic selection, HCC827 cells were injected into the flanks of nude mice. Injected mice developed subcutaneous flank tumors that were allowed to grow for 28 days, then were analyzed by targeted sequencing as described previously (3). We compared the normalized sgRNA representation among control non-targeting sgRNAs, sgRNAs targeting glycolytic and glycolysis-promoting genes, and sgRNAs targeting mito-ribosomal and respiratory chain (termed mito-respiratory) genes (Figure 1A). As a group, sgRNAs targeting mito-respiratory genes were significantly depleted compared to non-targeting control sgRNAs and sgRNAs targeting glycolytic genes. In fact, among all sgRNAs in the library, some individual sgRNAs targeting mito-respiratory genes, namely *MALSU1, HSD17β10, c14orf2*, and*TMEM261*, were the most significantly depleted individual sgRNAs (Figure 1A). Each of these genes is associated with mitochondrial function, and were previously found to be critical in maintaining mitochondrial-derived ATP levels, although none have known roles in tumorigenesis; *MALSU1* (Mitochondrial assembly of ribosomal large subunit 1) encodes a mitochondrial protein that is thought to be involved in mitochondrial translation (6). *HSD17β10* encodes Hydroxysteroid 17-Beta dehydrogenase 10, which localizes to the mitochondria and is involved in protein synthesis (7). *TMEM261* (also known as *DMAC1*) encodes transmembrane protein 261, an electron transport chain component (8). *c14orf2* (Chromosome 14 open reading frame 2, gene ATP5MPL) encodes ATP synthase membrane subunit j (9).

**Figure 1.**
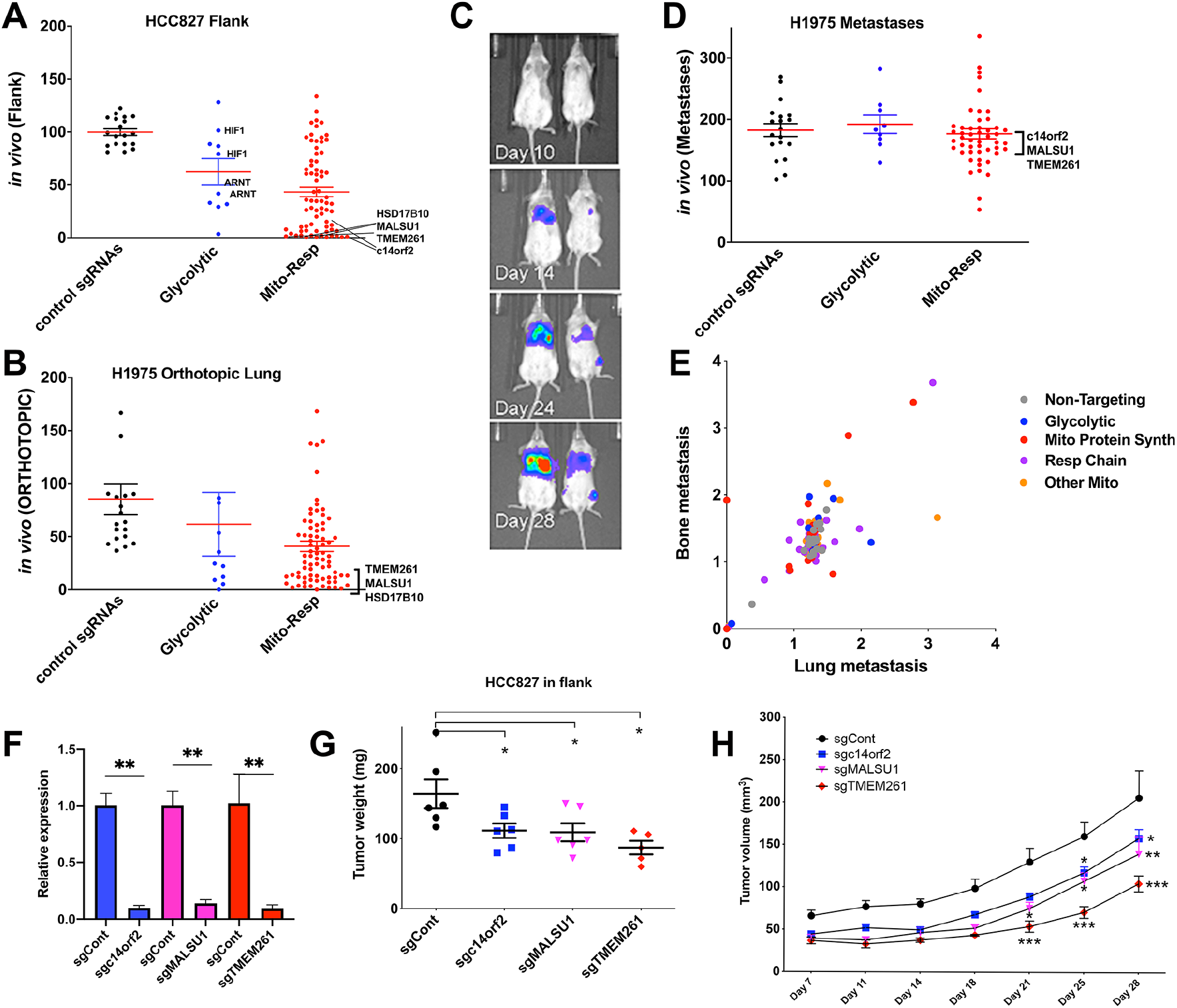
Human lung cancer cells demonstrate differential requirements for mitochondrial and respiratory genes when grown *in vivo* in flank or orthotopic lung models. HCC827 and H1975 human *EGFR*-mutant lung cancer cells were transduced with the mini-CRISPRi library and then were injected into either the subcutaneous space in flanks (HCC827) or into the left lower lung lobes (H1975) of nude mice and grown for 28 days. DNA from each tumor was sequenced and read counts for each sgRNA quantified. The read count for each sgRNA was normalized to the sum of reads for negative control sgRNAs and the ratio of each sgRNA’s frequency in the tumor model relative to its frequency *in vitro* (immediately pre-injection). A. HCC827 cells transduced with the CRISPRi mini-library were injected into the flanks of nude mice. Two independent replicate experiments were performed, with n=6 for each experiment (for n = 12 total). Each dot represents a single sgRNA and indicates its average normalized representation (computed from two independent replicate experiments), organized and displayed as control sgRNAs and sgRNAs targeting ATP-modulating genes classified as glycolytic or mito-respiratory (termed Mito-Resp). Mito-Resp sgRNAs were among the most severely depleted *in vivo* (*HSD17β10, TMEM261, MALSU1, c14orf2*) (mean and s.e.m. shown) (mean % representation of non-targeting, glycolytic and mito-respiratory sgRNAs as groups are 99.9%, 62.6% and 42.2% respectively. One-way ANOVA of all three groups of sgRNAs demonstrate p value <0.0001, with Tukey’s multiple comparisons test between glycolytic and control p<0.02, mito-resp and control p<0.0001, and between glycolytic and mito-resp ns). The complete list of sgRNAs comprising each class is available in Supplementary Table 1. B. H1975 human lung adenocarcinoma cells expressing Luciferase and the CRISPRi mini-library were injected into the left lungs of nude mice (n=6). H1975 cells grown orthotopically in the lungs of mice were dissected and analyzed using the same approach as for flank tumors (A) (mean % representation of non-targeting, glycolytic and mito-respiratory sgRNAs are 85.4%, 61.6% and 40.9% respectively. One-way ANOVA of all three groups of sgRNAs demonstrate p value = 0.006, with Tukey’s multiple comparisons test between glycolytic and control ns, mito-resp and control p-0.005, and between glycolytic and mito-resp ns). *In vivo* tumor growth of transduced H1975 cells injected into lungs and subsequent metastatic progression were monitored by bioluminescent imaging (C, two representative mice shown), which demonstrates tumor progression in the primary site, and metastatic spread to mediastinum and contralateral lung by day 28. In addition, an extra-thoracic distant metastasis (left femoral metastasis) developed (blue signal in the leg of the animal on the right). All metastases were confirmed at necropsy and analyzed by sequencing. D. Mean % representation of non-targeting, glycolytic and mito-respiratory sgRNAs in H1975 metastases (7 individual metastases from 4 mice, normalized to the matched primary tumor) are plotted on the y-axis (mean and s.e.m. shown, mean % representation of non-targeting, glycolytic and mito-respiratory sgRNAs are 183%, 192% and 177% respectively, one-way ANOVA of all three groups of sgRNAs with Tukey’s multiple comparisons test demonstrate ns). E. SgRNA frequencies in a lung metastasis is plotted on the x-axis against the identical sgRNA frequencies in the bone metastasis from the same animal. Dots representing individual sgRNAs are color coded by classification as non-targeting sgRNA, glycolytic, mitochondrial protein synthesis (Mito Protein Synth), respiratory chain (Resp Chain) and other mitochondrial function (Other Mito). F, G, H. Individually validating top mito-respiratory sgRNA indicates that silencing mito-translational genes suppresses *in vivo* tumor growth. Top mito-respiratory hits identified from the mini-CRISPRi library screen in HCC827 cells (Figure 1A) were individually cloned and expressed in HCC827 cells. sgRNAs targeting individual genes *c14orf2, MALSU1*, and *TMEM261* were transduced into HCC827 cells, selected with antibiotic, then injected into the flanks of nude mice. Tumors were allowed to grow for 28 days, then measured. F. qPCR analysis of transduced HCC827 cells assessing level of silencing achieved with single sgRNAs (mean and s.e.m. shown, t-test **p<0.01). G/H. Tumor weight (mg) and tumor volume (mm^3^) of HCC827 cells expressing each of the sgRNAs, mean and s.e.m. shown (Student’s t-test, *p<0.05, **p<0.01, ***p<0.001).

The depletion of individual sgRNAs in flank tumors after 28 days of *in vivo* growth may reflect cellular loss in response to metabolic pressures developing not only post-injection but at any time during the tumor growth period. We assessed early *in vivo* tumors for changes in sgRNA representation in a separate experiment in which HCC827 cells expressing the mini-library were grown as flank tumors for either four or seven days of growth (Supplementary Figure 1). Although control sgRNAs and glycolytic gene-targeting sgRNAs (∼95%) were comparably expressed across these timepoints, mito-translation and respiratory chain sgRNAs demonstrated significantly increased expression at day 4 (Supplementary Figure 1), before becoming significantly reduced in representation at day 7, preceding the near-complete depletion measured at day 28 (Figure 1A). These measured fluctuations in mito-respiratory sgRNAs at early timepoints in tumor establishment suggest metabolic adaptations involving these functions as tumors grow. The initial increased sgRNA representation (day 4) followed by reduction at day 7 may suggest an increasing reliance on mitochondrial function as tumors grow, perhaps due to an increased requirement for mitochondrial derived ATP as the tumor establishes itself.

To assess an alternate human tumor line and *in vivo* tumor model, we transduced the same mini-CRISPRi sgRNA library into H1975 human lung cancer cells expressing luciferase, which grow in both flank and orthotopic lung cancer models. In separate experiments, H1975 were injected in the flanks of nude mice or into the left lower lung lobes of C.B-17 SCID mice. Tumors in both models were grown for 28 days before dissection and analysis by sequencing.

Analysis of the sgRNA representation in H1975 tumors from the flank model demonstrated depletion of mito-respiratory sgRNA compared to control sgRNAs (Supplementary Figure 2), similar to our findings in HCC827 cells, although the difference in mean representation of the mito-respiratory sgRNAs and the level of significance were reduced compared to HCC827 cells grown in the flank.

As an alternative site of tumor growth, the orthotopic lung tumor model robustly produces intra-thoracic and distant metastases (Figure 1C). Tumors form at the primary site (orthotopically) then metastasize to mediastinal lymph nodes, the contralateral lung (Figure 1C) as well as distant sites, recapitulating the lethal pattern of disease spread in patients with lung cancer (10, 11). Both orthotopic and metastatic tumors were analyzed. Similar to flank tumors, tumors grown orthotopically (Figure 1B) also demonstrate significant reduction of mito-respiratory sgRNAs as a group and severe depletion of individual *MALSU1, HSD17β10, c14orf2, TMEM261* sgRNAs in primary lung tumors, as observed in HCC827 flank tumors. Orthotopic tumors in general develop in anatomically and physiologically distinct contexts that differ from subcutaneously grown flank tumors. However, these data show that primary tumors across cell lines and anatomic sites exhibit a strong requirement for mitochondrial function. Not only is this finding contrary to expectations, these data also specifically support a functionally significant need for mitochondrial-derived ATP, sharply contrasting with our findings in culture where respiratory chain genes were dispensable for growth (3).

### Mito-translational genes support *in vivo* tumor growth in primary sites but are dispensible in metastases

We then analyzed whether ATP-modulating genes correlate with metastatic tumor spread produced by injecting tumor cells orthotopically into the lung, which subsequently disseminated *in vivo* to produce regional (mediastinal), contralateral lung, and distant metastases, all of which were collected together with the primary tumor (10)(Fig. 1C). Interestingly, while sgRNAs targeting mitochondrial ribosomal and respiratory chain genes were severely depleted in orthotopic tumors (Fig. 1B), these same sgRNAs were enriched in metastases from the same animals (Figure 1D). As groups, control, glycolytic and mito-respiratory sgRNAs demonstrate comparable representation in metastases and no statistically significant differences were identified. These data suggest that ATP-modulating genes have differential effects in primary and metastatic tumors, as knocking down mitochondrial genes that are essential for primary tumor growth was dispensable in metastatic tumor deposits in the same mouse model. Each mouse developed on average two separate metastatic deposits and individual metastases from a given mouse were compared. Interestingly, we found that sgRNA representation between anatomically discrete metastases from the same mouse correlated across functional classes (non-targeting, glycolytic, respiratory chain, mitochondrial protein synthesis). We analyzed metastases from a mouse that developed both lung and leg bone metastases (the only extra-thoracic metastasis) and found significant correlation in sgRNA representation between most sgRNA classes (Figure 1E), suggesting generalizability of mito-respiratory gene effects to more than one metastatic context.

### Expressing individual sgRNAs against mito-respiratory hits suppresses *in vivo* tumor growth

*c14orf2, MALSU1*, and *TMEM261* encode mitochondrial proteins that were previously identified as strong low ATP hits under respiratory conditions (3); these genes are among the strongest growth repressive hits in both the flank and orthotopic tumor growth screen. CRISPRi sgRNAs targeting each of these individual genes *c14orf2, MALSU1*, and *TMEM261* were expressed in HCC827 cells, and after silencing was confirmed (Figure 1F), these cells were injected into the flanks of nude mice. Tumors were grown for 28 days, then measured immediately after removal (Figure 1 G/H). Each of the CRISPRi sgRNAs targeting mito-respiratory hits significantly decreased *in vivo* tumor growth in the flank compared to pseudogene control sgRNA. *C14orf2, MALSU1*, and *TMEM261* have no known roles in supporting tumor growth, however these data provide evidence that their functions promote tumor growth and collectively suggest that mitochondrial genes are necessary for *in vivo* tumor growth. We assessed the tumors for reduced mitochondrial content as a possible consequence of silencing and cause of reduced respiratory function by performing Western blotting for the mitochondrial protein TOMM20. All tumors from each sgRNA-expressing cell line demonstrated comparable TOMM20 protein levels across all sgRNAs (Supplementary Figure 3), consistent with maintained mitochondrial content in the context of the silenced mito-respiratory hits.

### Suppression of mito-respiratory hits is associated with overexpression of glycolytic genes *in vitro* and silencing of glycolytic genes *in vivo*

To gain insight into the convergent mechanisms by which these genes impact respiration, we performed RNA Seq and transcriptional profiling analysis of HCC827 cells expressing sgRNAs against *MALSU1, c14orf2* or *TMEM261* compared to control sgRNA-expressing HCC827 cells grown under basal conditions (Fig. 2A). All three mito-respiratory sgRNAs are associated with significant overexpression of glycolytic genes (Fig. 2A), with *PGK1, ENO2* and *HK2* being the most significantly overexpressed glycolytic genes in all three analyzed lines (Fig. 2A/B). In addition, gene set enrichment analysis identified glycolysis as being most significantly altered among all three cell lines (Fig. 2B). These concordant data support transcriptional upregulation of glycolytic genes as a shared compensatory mechanism utilized by cancer cells when mito-respiratory function is handicapped in order to enable increased glycolytic flux.

**Figure 2.**
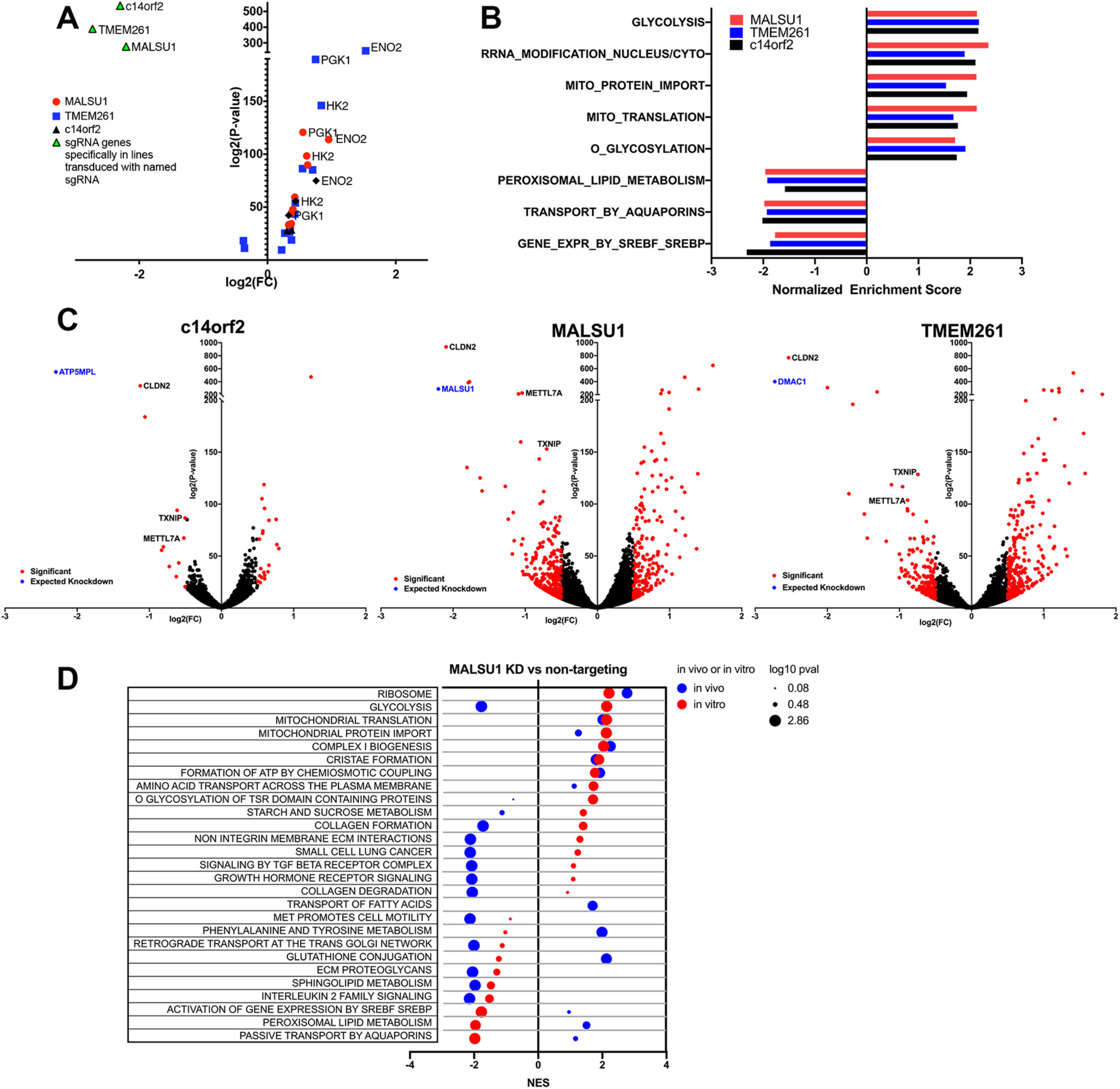
Transcriptome profiling of human lung cancer cells expressing sgRNAs against mito-respiratory hits reveals overexpression of glycolytic pathway genes *in vitro* and underexpression *in vivo*. RNA-Seq was performed on HCC827 cells expressing individual sgRNA against top hits *c14orf2, MALSU1, TMEM261* (in addition to control sgRNA) (n=4 samples per cell line) were analyzed for changes in gene expression *in vitro* and *in vivo*. A. Averaged expression for each mito-respiratory sgRNA was compared to control cells, fold-change, indicated by log2(FC) plotted on the x-axis, and log2(P-value) on the y-axis. SgRNA-targeted genes are indicated by green triangles, with each cell line demonstrating expected significantly reduced expression of its targeted gene (upper left quadrant). Glycolytic genes were the most overexpressed genes overall, with *ENO2, PGK1* and *HK2* being the most over-expressed genes across all three cell lines. B. Gene set enrichment analysis indicate glycolysis and mitochondria-associated pathways and ontologies as enriched in all three cell lines. Significance was set at p<0.05. C. Volcano plots shown for each of the single sgRNA cell lines. Expected sgRNA-mediated silencing was observed in all three cell lines (blue dot). D. HCC827 cells transduced with either control sgRNA or sgRNA MALSU1 were injected into the flanks of nude mice, then grown for 28 days, after which tumors were removed and analyzed by RNA Seq. Pathway analysis was performed comparing the sgRNA MALSU1 cells to control sgRNA cells grown *in vivo* (blue) or *in vitro* (red), and displayed in a bubble plot indicating normalized enrichment score (NES) and log_10_ p-value for significantly altered pathways. Control sgRNA tumors n = 4, sgMALSU1 tumors n = 2. All source data are provided as Supplementary Source Data.

Transcriptome profiling indicated that reactive oxygen species, glutathione and NADPH, mitochondrial functions not directly involved in ATP production, were not consistently altered in the context of mito-respiratory gene silencing. In contrast, the expression of genes involved in fatty acid β oxidation was significantly reduced in cells in which *MALSU1, c14orf2* or *TMEM261* is silenced compared to control sgRNA-expressing HCC827 cells (Supplementary Table 1). SREBP-regulated gene expression, which regulates lipogenesis as well as growth and mitochondrial metabolism in some cancer cells, was also decreased with all three mitochondrial genes, suggesting cross-talk between mitochondrial function and lipogenesis. Outside of the broad expression changes in major pathways, individual cell lines also demonstrated shared significantly decreased expression of *CLDN2, METTL7A* and *TXNIP* (Fig. 2C). *CLDN2* encodes claudin-2, a tight junction protein (12). *METTL17* encodes a mitochondrial protein involved in the translation of mitochondrial coding genes (13). *TXNIP*, a thioredoxin-binding protein involved in redox regulation, and functioning as a tumor suppressor gene (14). None of these genes are known to interact with each other, and thus their downregulation in each of the mito-respiratory silenced cell lines implicates all three as participating in a transcriptional response triggered by decreased mitochondrial ATP levels. Other non-ATP functions were transcriptionally altered only in selected cell lines; i.e., transcripts for genes involved in glutathione metabolism were significantly reduced in *MALSU1* cells.

HCC827 cells transduced with either control sgRNA or sgRNA *MALSU1* were injected into the flanks of nude mice, then grown for 28 days, after which tumors were removed and analyzed by RNA Seq. Pathway analysis was performed to compare sgRNA control cells grown *in vivo* to the same cells grown *in vitro* (Supplementary Figure 4). Tumor cells grown *in vivo* demonstrated increased expression of genes involved in interferon signaling, collagen degradation, collagen chain trimerization and ECM (Supplementary Figure 4A), consistent with the involvement of these processes in *in vivo* tumor growth. Comparing pathway enrichment for sgRNA *MALSU1* cells to control sgRNA cells when grown either *in vivo* or *in vitro* demonstrates significant similarities that are likely *MALSU1*-loss associated, notably in ribosome, mitochondrial translation, mitochondrial protein import, complex I biogenesis and cristae formation (Figure 2D, Supplementary Figure 4B).

Our analysis also identified significant differences in transcriptomic response with *MALSU1*-silencing that are context-dependent; expression of glycolysis pathway genes, while increased *in vitro*, is significantly reduced in sgRNA *MALSU1* cells grown *in vivo* (Figure 2D), supportive of the concept that tumor cells grown *in vivo* instead optimize respiratory function (3). These differences indicate that the *in vivo* tumor growth context accentuates the significance of some pathways, and given that mitochondrial and respiratory-driven ATP is substrate-dependent, suggests that these differences in transcriptome profiles reflect tumor responses to substrate restriction.

### Silencing Mito-respiratory genes suppresses TCA cycle activity

Compensatory shifts in gene expression (from respiration to glycolysis and vice versa) occur in a context-dependent manner after silencing mito-respiratory hits (Figure 2). To determine how mito-respiratory hits *c14orf2, TMEM261*, and *MALSU1* alter metabolite levels and the pathways in which they function we performed ^13^C-glucose and ^13^C-glutamine-based metabolomics analysis of HCC827 cells expressing individual CRISPRi sgRNAs against these genes. HCC827 cells expressing individual CRISPRi sgRNA (control, or mito-respiratory hits *TMEM261, MALSU1*, or *c14orf2*), were grown under either basal or respiratory (10 mM 2DG), or glycolytic (oligomycin) conditions with either ^13^C-glucose or ^13^C-glutamine for 18 hours. Cells were then collected, metabolites extracted and analyzed by mass spectroscopy.

Metabolomics analysis demonstrates the relative differences in sources of TCA cycle metabolites, with Gln being the predominant source (Supplementary Fig. 5). Glc labeling resulted in relatively low percent labeling of citrate (ranging from 5 – 30%, seen in Supplementary Fig. 5A right panel), whereas Gln labeling of most TCA cycle metabolites was approximately 80% or greater (Supplementary Fig. 5C, right panel). Overall, total amounts of TCA cycle metabolites varied between the four cell lines, with Glc and Gln labeling demonstrating similar differences between cell lines under both basal and 2DG conditions (Supplementary Fig. 5). Basal growth of *TMEM261* and *MALSU1*-silenced cells is associated with significant reduction in the percent labeling of the TCA metabolites citrate and aconitate (Supplementary Figure 5A), while sg*c14orf2* cells resemble control cells. Growth under glycolytic block (2DG) generally decreased ^13^C incorporated into these TCA metabolites when compared to basal conditions, as expected, but here also sgc14orf2 cells showed reduced labeling of citrate and aconitate compared to control cells. Downstream in the TCA cycle with the addition of 2DG, *c14orf2* silencing still results in a reduction of Glc-labeling of citrate, similar to *TMEM261* and *MALSU1*, indicating a deficiency shared by all three mito-respiratory-hits (Supplementary Fig. 5B, right panel); since this effect is only visible with a glycolytic block, *c14orf2* may be more dispensable than either *TMEM261* and *MALSUI* when glucose is available. ^13^C-glutamine-labeling also distinguished metabolite labeling profiles for cells in which mito-respiratory hits were silenced. In comparison to labeling under basal control conditions, under forced respiration (2DG, Supplementary Fig. 5C/D, right panels) ^13^C incorporation into succinate was the most significantly reduced in all three mito-respiratory lines compared to control.

### Silencing Mito-respiratory hits shifts cells to greater glycolytic metabolism *in vitro*

Genetically-mediated modulation of cellular ATP involves genes that concurrently optimize one metabolic pathway while suppressing the alternative pathway (3). To detect metabolic shifts towards glycolytic or respiratory function, we compared isotopologues of glycolytic and respiratory metabolites among mito-respiratory-deficient cells, grown under basal or forced respiratory (2DG) conditions. Examining as an index glycolytic metabolite fructose 1, 6-bisphosphate (F16BP), under basal conditions, the unlabeled (0 carbon) F16BP in sgRNA *MALSU1* and sgRNA *c14orf2* cells is comparable to control and modestly decreased in sgRNA *TMEM261* cells, while fully labeled (6 carbon) F16BP is comparable between control and *TMEM261* as well as *c14orf2* cells, and modestly reduced in *MALSU1* cells (Supplementary Fig. 7A, upper right and left panels). To summarize, under basal metabolic conditions F16BP in mito-respiratory silenced cells resemble the same metabolite in control sgRNA cells.

Adding 2DG however reveals glycolytic metabolism in mito-respiratory cells similarly diverging from control cells (Supplementary Fig. 7A, lower right and left panels). In the presence of 2DG, all three mito-respiratory silenced cell lines continue labeling the downstream glycolytic metabolite F16BP (6 carbon labeled), in contrast to control cells (Supplementary Fig. 7A, lower right panel), consistent with increased relative glycolytic capacity compared to control cells.

We similarly compared Gln-labeled glutamate among the cell lines under basal and forced respiratory conditions (Supplementary Fig. 7B). Under basal growth, sgRNA *TMEM261*, sgRNA *c14orf2* and sgRNA *MALSU1* cells all demonstrate a marked increase in the percentage of unlabeled glutamate compared to control cells (Supplementary Fig. 7B). Fully labeled glutamate (5 carbons labeled) distinguishes *c14orf2*-silenced cells as demonstrating reduced labeling compared to *TMEM261* and *MALSU1*-silenced cells (Supplementary Fig. 7B right panel). These differences in Gln-derived labeling among mito-respiratory hits presumably reflects the discrete defects in respiratory metabolism associated with each of the mito-respiratory hits, which suggests more defect-specific metabolic signatures detectable by Gln-labeling.

Quantifying the incorporation of Glc and Gln-labeled metabolites provide approximations of the utilization of glycolysis and respiration, and combining the unlabeled and labeled F16BP and glutamate quantifications as ratios of glycolytic label incorporation (6C/unlabeled F16BP) to the respiratory label incorporation (5C/unlabeled glutamate) for *TMEM261, c14orf2* and *MALSU1* knockdown indicate glycolysis-shifted metabolic metabolism in the three mito-respiratory silenced cell lines (Figure 3C) and a shared metabolic response amongst these three genes. Thus, while the absolute magnitude of basal Glc and/or Gln-labeling in individual glycolytic and respiratory metabolites may vary between each of the mito-respiratory sgRNAs as compared to control cells, the relative incorporation patterns from combined Glc and Gln-labeling highlight increased glycolytic utilization with *TMEM261, c14orf2* and *MALSU1* knockdown.

**Fig. 3.**
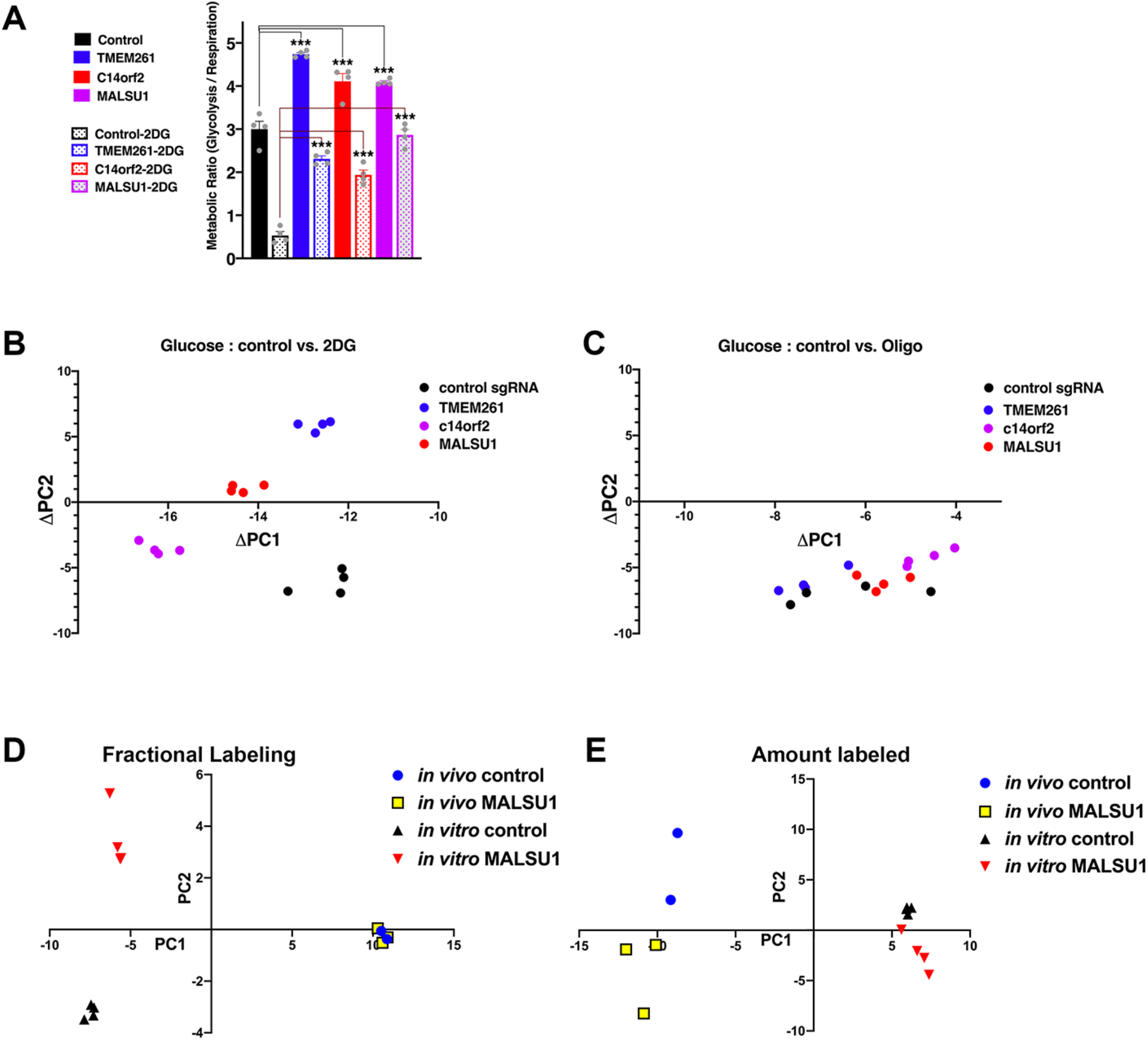
Mito-respiratory hits have distinct metabolic signatures that differ between *in vitro* and *in vivo* contexts. HCC827 cells expressing individual CRISPRi sgRNA (control, or mito-respiratory hits *TMEM261, c14orf2, MALSU1*, silencing confirmation shown in Fig. 2A), were grown in basal media or media with 10 mM 2DG (n=4 per group, individual data points shown as grey dots) with either [U-^13^C]glucose or [U-^13^C]glutamine for 18 hours. Cells were collected, metabolites extracted and analyzed by mass spectrometry. **A**. Using the measured % of total all labeled and unlabeled glycolytic [[U-^13^C] glucose→ fructose 1,6-bisphosphate indicates % total unlabeled (0 carbon) or completely labeled (6 carbon)] and respiratory metabolite values [[U-^13^C]glutamine→ glutamate analysis indicates total unlabeled (0 carbon labeled) on the left and fully labeled (all 5 carbons labeled)](shown in Supplementary Figure 6A/B), the estimated metabolic flux ratios of glycolytic flux (6C/unlabeled) to the respiratory flux (5C/unlabeled) are shown for 3 mito-respiratory hits (*TMEM261, c14orf2* and *MALSU1*). The ratios distinguish the glycolysis-shifted metabolism apparent in the three cell lines in which mito-respiratory hits are silenced. (n=4 replicates per sample, 1-way ANOVA, Dunnett’s multiple comparisons test, *p<0.05, ***p<0.001). **B/C**. Principal component analysis was applied to the fractional contribution values of the metabolomics data for HCC827 cells expressing individual CRISPRi sgRNA (control, or mito-respiratory hits *TMEM261, MALSU1*, or *c14orf2*). ^13^C glucose-derived labeling of cells grown under either control or 2DG media (D) was compared in **B**, control or oligomycin in **C**. The absolute change in PC1 and PC2 values of the fractional contribution analysis for each cell line comparing control to 2DG growth (ΔPC1 = PC1_control_ – PC1_2DG_, ΔPC2 = PC2_control_ – PC2_2DG_) shown in **B** or control to oligomycin shown in **C** are plotted (n=4 replicates). **D/E**. HCC827 cells expressing either control or *MALSU1* sgRNA were injected into the flanks of nude mice. After 28 days of growth, mice were injected with ^13^C glucose to label tumor metabolites, after which tumors were collected and metabolites analyzed. **D**. Using the fractional labeling values of glucose-derived metabolites we performed PCA and compared *MALSU1*-deficient and control HCC827 cells grown *in vitro* or *in vivo*. **E**. PCA was performed on the amount labeled of glucose-derived metabolites in MALSU1-deficient and control HCC827 cells grown *in vitro* or *in vivo*.

### Silencing of ATP-modulating mito-respiratory genes is associated with discrete metabolite profiles and metabolic network structure that distinguish *in vitro* and *in vivo* growth

We then examined metabolic networks on a broader scale by performing principal component analysis (PCA) of the metabolomics data (Figure 3 and Supplementary Figure 8). After computing Principal Component (PC)1 and PC2 scores for control and mito-respiratory silenced HCC827 tumor cells that were grown under either control vs. 2DG, or control vs. oligomycin (Supplementary Figure 5), we plotted the ΔPC1:ΔPC2 score between control and 2DG for each cell line (Figure 3B) to depict the net magnitude and directionality of PC score changes between control and 2DG, which fail to support a common respiration-driven metabolite signature among all the mito-respiratory gene hit silencings. However, in contrast to 2DG treatment, oligomycin treatment (forced glycolysis) was associated with similar net shifts in PC1 and PC2 scores among the four cellular genotypes (Figure 3C), suggesting a common glycolytic response for all mito-respiratory hits and contrasting with the divergent respiratory response in these same cells (Figure 3B).

Considering the differential requirement for mito-respiratory hits *in vitro* versus *in vivo*, we next assessed metabolites *in vivo*. We injected HCC827 cells expressing either control or *MALSU1* sgRNA into the flanks of nude mice, grew tumors for 28 days, then injected mice with ^13^C glucose, after which tumors were collected and metabolites analyzed. We performed PCA based on either the fractional contributions or the amounts of labeled glucose to measured metabolites (Figure 3D/E). These comparative analyses show clear growth context-dependence (*in vitro* vs. *in vivo*) of metabolites in general, however also indicate *MALSU1*-specific effects that are distinguishable in a growth context-dependent manner. PC1 describes most of the variation between metabolite pool sizes and fractional labeling (62.06% and 75.52% of the variation, respectively) *in vitro* vs *in vivo*, which may result from differences in substrate introduction and metabolite extraction, however, PC2 describe variation in metabolite pool sizes and fractional labeling (16.42% and 7.7% of the variation, respectively) not associated with *in vitro* vs *in vivo* difference, likely attributable to *MALSU1* knockdown. On the basis of fractional contributions, samples from *MALSU1* knockdown cells and control cells clustered *in vivo* but separated *in vitro* (Figure 3D). Conversely, PCA based on metabolite amounts separates samples from *MALSU1* knockdown cells and samples from control cells *in vivo* but marginally *in vitro* (Figure 3E), suggesting that *MALSU1*-silencing effects on the metabolic network *in vivo* are driven by discrete metabolites.

We then sought to assess metabolic network structure for each of the mito-respiratory genes by integrating transcriptomic and metabolomic data on metabolism pathway-focused graphs (15-17)(Figure 4). All three mito-respiratory-deficient cells shared similar activation of nodes in glycolytic metabolism (Figure 4A-C) and similar reduction in TCA cycle network (Supplementary Figure 8); these shared features suggest a conserved metabolic network structure occurring among mito-respiratory functional deficits. Furthermore, similar integrated transcriptomic and metabolomic-based pathway analysis of glucose-labeling by *MALSU1*-deficient cells grown *in vivo* revealed a striking reduction in glycolytic metabolite pools as compared to *in vitro* growth (Figure 4D), consistent with the transcriptional changes summarized in Figure 2D. These data provide further evidence that mito-respiratory-deficient cells engage an *in vivo* context-dependent metabolic network that is distinguished by glycolytic function (Figure 4E), and these differences may underlie the differential substrate-driven requirement of these genes *in vivo* and *in vitro*.

**Figure 4.**
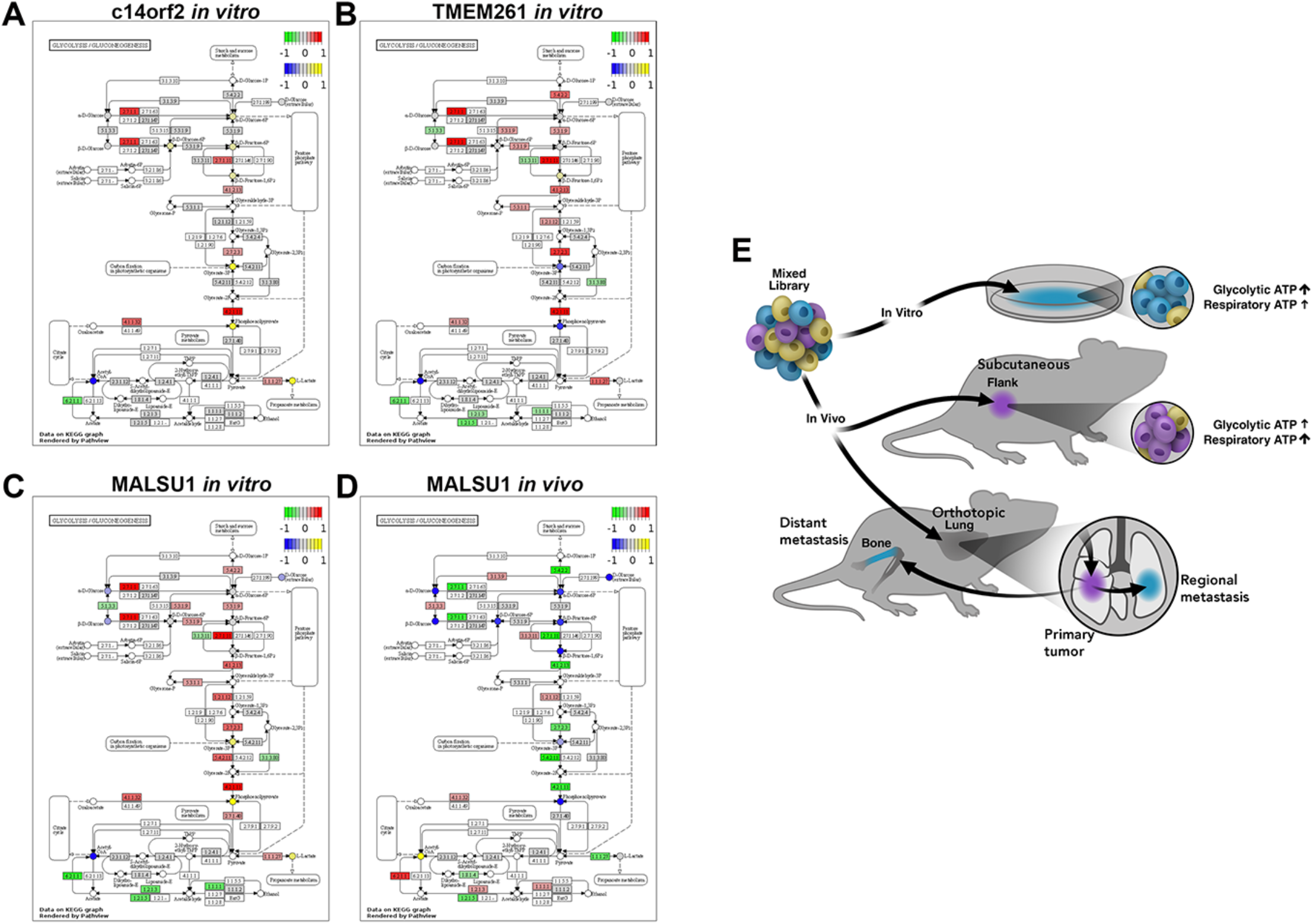
Integrated principal component and pathway analysis of metabolic network structure indicates differential effects of mito-respiratory hits *in vitro* and *in vivo*. **A/B/C**. HCC827 cells expressing individual CRISPRi sgRNA (control or mito-respiratory hits *c14orf2, TMEM261*, or *MALSU1*) were grown *in vitro* and analyzed by RNA-Seq and metabolomics (described in Figures 2 and 3). Pathway analysis was performed by integrating transcriptome and metabolomics data and plotted on a KEGG graph depicting pathway components for glycolysis/gluconeogenesis. **D**. Pathway analysis was performed by integrating transcriptome and metabolomics data for HCC827 cells expressing individual CRISPRi sgRNA (control or *MALSU1*) grown *in vivo* as flank tumors. **E**. Systems-level testing implicates context-specific growth effects of mitochondrial/respiratory function and ATP levels. ATP-modulating CRISPRi hits grown in lung cancer cells produce specific energy substrate-driven growth effects *in vitro* and *in vivo. In vitro* growth correlates with glycolytic ATP, while growth in the subcutaneous and orthotopic lung setting correlate with mitochondrial-derived ATP.

## DISCUSSION

Previous studies have implicated cell-intrinsic metabolic differences as important determinants of metastatic potential (18). However, primary tumors and subsequent metastases occupy diverse anatomic locations and growth in different sites may require alternate metabolic processes specific to the local environment (19). These locally-defined requirements and what metabolic capabilities tumor cells must possess to negotiate these remain to be well-defined and therapeutically targetable, but would have significant clinical implications if understood. Metastatic disease remains essentially fatal and causes serious morbidity in cancer patients, and therefore improving the control of any subset of metastases would likely be positively impactful for patients.

Lung cancers in particular are known to utilize multiple metabolic pathways to support growth *in vitro* (20), however which metabolic pathways are required for *in vivo* growth of primary and metastatic lung cancer is not well understood. In our study, human lung cancer cells were grown in multiple *in vivo* contexts to test whether genes modulating ATP were selectively enriched in cells grown as solid tumors in each of these contexts. The enrichment and depletion of select classes of genes implicates specific metabolic requirements for *in vivo* tumor growth. These contrasts in functional outcome between *in vitro* and *in vivo* tumor growth suggest that tumor cells utilize aerobic respiration in a context-dependent manner to meet ATP requirements *in vivo* (Fig. 4E), rather than being hard-wired to preferentially utilize aerobic glycolysis at all times.

*c14orf2, MALSU1*, and *TMEM261* were top CRISPRi ATP-modulating hits that each underlie respiratory chain dysfunction, severely impacting ATP production (3) as well as *in vivo* tumor growth. The mechanisms by which each of these genes accomplishes this is not fully known; we previously found that silencing of another respiratory ATP-modulating gene HSD17β10 was associated with a relatively slow rate of ATP depletion under respiratory metabolism but relatively rapid rate of ATP depletion under basal substrate conditions (3). This example indicates that mechanisms promoting either ATP production or ATP consumption can each contribute to ATP levels and metabolic phenotype in disease-defining ways. *c14orf2*, which is part of ATP synthase/complex V, *MALSU1* (apparently needed for normal translation or function of mitochondrial ribosomes), and *TMEM261* (a component of complex I) are mitochondrial proteins, although there are no known functional connections between these genes and how they act on respiration.

Both metabolomics analysis and transcriptional profiling analysis of the three mito-respiratory hits (Fig. 3) identified a unified transcriptional and metabolomic signature that notably shifts glycolytic utilization. Importantly, expression of glycolysis pathway genes differentiated on the basis of environmental context, being significantly reduced *in vivo*, but not *in vitro. In vitro*, tumor cells boost glycolysis to compensate for impaired respiration, which enables them to survive and explains their distinct transcriptional/metabolomic profile. This shift to glycolysis is specific to the context in which tumor cells grow; both the transcriptomes and metabolomes of respiratory ATP-deficient cells grown *in vitro* versus *in vivo* demonstrated starkly contrasting glycolytic responses, only in the former case being consistent with the Warburg effect.

The *in vivo* growth differences based on gene function may involve metabolic plasticity on the part of tumor cells to successfully colonize different anatomic sites and metabolic environments. However, tumor cells also interact with their environments; analysis of tumor microenvironments by tumor interstitial fluid analysis, for example, have quantified differences in metabolite levels between tumor locations (19). Our data suggest that tumor cells shift to glycolytic metabolism when disseminating and establishing distant tumor deposits, contrasting with primary tumor growth at an initial site of origin, perhaps to support increased biosynthesis needed for the process of establishing metastases and/or in response to limited nutrients/oxygen.

Other contributing mechanisms of metastasis, such as cellular migration and invasion, escape from immune surveillance, and others, could invoke alternative metabolic adaptations (such as production of specific metabolites and alternate bioenergetic processes). One limitation of our study is that we analyzed established *in vivo* metastases that had clearly completed multiple steps of metastasis formation. The scope of this initial work does not enable us to resolve the intermediate steps of metastasis formation and thus the dependencies we observe may localize to different stages of metastases. An obvious question would also be how cancer cells transition between these points and elucidating what may be metabolically intermediate states through which metastases progress. Indeed, our own data strongly hint at dynamic alterations, as mito-respiratory sgRNAs fluctuated significantly increased between initial injection and day 4 of flank growth, before significantly depleting between day 4 and day 7 (Supplementary Figure 1). Elucidating these steps will require appropriately tailored modeling approaches.

Our study focused on lung cancers and it should be acknowledged that other tumor cell types as well as other anatomic sites of origin may shift differently between glycolytic and respiratory metabolism, invoking other metabolic processes. Melanoma tumor cell survival and *in vivo* disease progression, including metastasis, have been shown to be facilitated by suppression of ferroptosis (21, 22). The potential diversity of metabolic phenotypes assumed by tumor cells should be experimentally elaborated and may contribute to the general goal of eradicating metastatic disease.

Metastatic disease is genetically polyclonal, and by taking a systems-driven experimental approach into physiologic contexts our work contributes to understanding metastases as poly-metabolic, that is, involving discrete metabolic states that are likely adapted to the local metabolism and a cell’s bioenergetic needs (in fact our CRISPRi screening library is comprised of ATP-modulating hits from a previously published genome-wide ATP screen (3)) and thus may also point to specific vulnerabilities. To date, metabolism-based therapies are not routinely integrated into cancer management. Greater basic understanding of cancer metabolism to inform metabolically-based treatment approaches is needed and could comprise a productive strategy.

## MATERIALS AND METHODS

### Cell culture

The human lung adenocarcinoma HCC827 cell line was originally obtained from Trever Bivona (UCSF) (ATCC 2868, 39-year-old Caucasian female individual). HCC827 cells were grown at 37 **°**C in RPMI medium with 10% FBS, 1% penicillin/streptomycin. H1975 cells expressing GFP-luciferase were grown at 37 **°**C in DMEM medium with 10% FBS, 1% penicillin/streptomycin.

### RNA isolation, reverse transcription (RT), and real-time RT-PCR to confirm gene knock-down

RNeasy Mini Kit (Qiagen) was used to isolate total RNA. SuperScript IV (ThermoFisher Scientific) was used to synthesize cDNA. Gene expression was measured by Real-time PCR on QuantStudio 5 using Taqman probes. All of Taqman probes were purchased from ThermoFisher Scientific, and assay IDs are Beta-Actin (Hs99999903_m1), HSD17B10 (Hs00189576_m1), MALSU1 (Hs00370770_m1), TMEM261 (Hs00383923_m1), and c14orf2 (Hs01043634_m1). cDNAs and PCR reactions were prepared according to the protocol for Cells-to-CT kit (ThermoFisher #AM1728), using the standard reverse transcription cycle (95°C for 2 min, inactivation at 95°C for 5 min, hold at 4°C), and qRT-PCR conditions (UDG Incubation – 50°C for 2 min, enzyme activation – 95°C for 10 min, PCR cycle – 95°C for 15 sec, 60°C for 1 min – repeat 40 cycles). All reactions were performed in a 384-well plate, in replicates of at least n=3 and from 2 independent experiments. CT (threshold cycle) values of each gene were averaged and calculated relatively to CT values of β-actin using the 2^−△△CT^ method^(23)^.

### Flank xenografts

Surgical procedures and all animal work was done under a protocol AN182206-02, approved by the UCSF Institutional Animal Care and Use Committee (IACUC). Transduced cells were injected (1 × 10^6^ cells) into the right flank of nude mice as previously described(24). Injected mice developed subcutaneous flank tumors that grew for 28 days then were analyzed by sequencing.

### Orthotopic lung xenografts in immunodeficient mice

Six to eight week old female SCID CB.17 mice were obtained from Charles River and housed in pathogen-free conditions and facilities as previously described (10). Briefly, tumor cells expressing GFP-Luc were suspended in Matrigel (Corning). Cell concentration was adjusted to 1 × 10^5^ cells/ul, and cell suspension was transferred into a 1 ml syringe. Syringe was kept on ice until implantation. After anesthetizing, 1 cm surgical incision was made along the posterior medial line of the left thorax. 10 ul cell suspension were injected into left lung directly. Visorb 4.0 polyglycolic acid sutures were used for primary wound closure of the skin layer.

### Bioluminescence imaging

Xenogen IVIS-100 (PerkinElmer) were used for in vivo imaging. Mice were injected intraperitoneally with VivoGlo vivoGLO luciferin, *in vivo* grade : P1042(Promega). Mice were injected intraperitoneally with vivoGLO luciferin (150mg/kg). After 10 minutes, mice were anesthetized with 2% isoflurane, and then they were transferred into Xenogen IVIS-100. All mice were imaged twice weekly. Living Image (PerkinElmer) was used for analysis. All of imaging process followed manufacturer’s instruction.

### In Vitro Metabolomics

5 × 10^5^ Cells per well were plated in 6 well plates. After 24 hours, cell medium were aspirated. Cells were rinsed quickly with ice-cold 150mM NH4AcO (pH7.3). After removing NH4AcO, added 1ml 80% MeOH which were precooled on dry ice. Plates were incubated on dry ice for 20 minutes. Each samples were transferred into 1.5ml tube placed on ice, and vortex each tubes for 10 seconds. After centrifuge at 16,000g for 15min at 4°C, supernatant were transferred into new tubes. Samples were dried up in a speed vac. Samples were stored in -80°C. All samples were analyzed by the UCLA metabolomics center (3).

HCC827 cells were incubated for 2 and 6 h in either (1) respiratory condition: 2% FBS, 10 mM [U-^13^C]pyruvate + 10 mM 2DG, (2) glycolytic condition: 2% FBS, 2 mM [U-^13^C]glucose + 5 μM oligomycin + 3 mM 2-deoxyglycose, (3) basal condition: 2% FBS, 10 mM [U-^13^C]glucose + 5 mM pyr-uvate with no drugs, or (4) 2% FBS, 10 mM [U-^13^C]pyruvate alone, before metabolite extraction in 80% methanol and drying in a Labconco CentriVap. Dried metabolites were resuspended in 50% ACN:water and loaded onto a Luna 3um NH2 100 A (150 × 2.0 mm) column (Phenomenex). The chromatographic separation was performed on a Vanquish Flex (Thermo Scientific) with mobile phases A (5 mM NH4AcO, pH 9.9) and B (ACN) and a flow rate of 200 μL/min. A linear gradient from 15% A to 95% A over 18 min was followed by 9 min isocratic flow at 95% A and re-equilibration to 15% A. Metabolites were detection with a Q Exactive mass spectrometer (Thermo Scientific) run with polarity switching (+3.5 kV/−3.5 kV) in full scan mode with an m/z range of 65– 975. TraceFinder 4.1 (Thermo Scientific) was used to quantify the targeted metabolites by area under the curve, using expected retention time and accurate mass measurements (<5 p.p.m.). Values were normalized to cell number. Relative amounts of metabolites were calculated by summing up the values for all isotopologues of a given metabolite. Metabolite isotopologue distributions were corrected for natural C13 abundance.

### In Vivo Metabolomics

Mice undergoing the procedure were fasted for 12 hours prior to the infusion of ^13^C-glucose. The mice were weighed the morning of the procedure and anesthetized with 1.5% isoflurane (v/v). Catheters constructed from 30 g needles (BD 305106) and polyethylene tubing (BD 427400) were used to infuse a 200 µL bolus of ^13^C-glucose (0.4 mg/g) into the tail vein. Following the bolus, a 150 µL/h infusion of ^13^C-glucose at a dosage of 0.012 mg/g/min was administered for 30 min (25-27). Mice were then euthanized and the tumors were flash frozen in an isopentane bath cooled to -80ºC with dry ice. To extract the metabolites from the frozen tissue, the samples were first homogenized in a cryogenic mortar and pestle before being mixed with 1 mL of 80% methanol chilled to -80ºC. Samples were then vortexed for 20 s and incubated at -80ºC for 20 minutes. Following the incubation, the samples were vortexed for an additional 20 s and then centrifuged at 16,000 g for 15 minutes in a 4ºC chamber. The supernatant was transferred to a -80ºC pre-chilled 1.5 mL tube. To normalize the extracted metabolite amounts by protein content, a BCA assay was completed on the sample pellets. A 100 µg protein equivalent of extracted metabolite was aliquoted from each sample and dried in a Labconco CentriVap. The dried samples were then stored at -80ºC and analyzed by the UCLA Metabolomics Center.

### Western Blotting

Western blotting was performed as previously described (28) using TOMM20 antibody(ab186735)(Abcam).

### Sequencing and computation of sgRNA representation

Ortho: in vitro - divides each primary tumor by the average for in vitro (n=6) in denominator dataset, remove sgRNAs that have <200 reads (200 is the threshold) <50 was for ATP-sensor FRET FACs)

Genomic DNA was isolated using the Macherey-Nagel NucleoBond Xtra Midi Plus (Macherey-Nagel, Germany). The sgRNAs were amplified and adaptors attached in a single PCR step. 1.5 µg of undigested genomic DNA was used per 50 µL PCR reaction, and sufficient reactions were performed to include all isolated genomic DNA. PCR was conducted using Q5 HotStart High Fidelity Polymerase (NEB, Ipswich, MA) using forward primer: aatgatacggcgaccaccgaGATCGGAAGAGCACACGTCTGAACTCCAGTCACNNNNNNgcacaaaaggaaactcac cct 1and reverse primer: caagcagaagacggcatacgaCGACTCGGTGCCACTTTTTC, which include necessary adaptor and indexing sequences. “N” refers to the variable index sequence. PCR parameters were 98°C for 30 sec, followed by 26 cycles of 98°C for 15 sec, 62.5°C for 15 sec, 72°C for 20 sec and ending with 72°C for 6 min; samples were then ramped down to 4°C and held. The resulting PCR product from multiple reactions were pooled, and unincorporated primers were removed using the GeneRead Size Selection Kit. Quality and purity of the PCR product was assessed by bioanalyzer (Agilent) and sequencing was performed on an Illumina HiSeq 2500 as described (29). Informatic analysis of the raw reads was performed as described (29).

### Quantification and Statistical Analysis

All statistical analyses, including the n, what n represents, description of error bars, statistical tests used and level of significance, are stated in the figure legends. All measurements were taken from distinct samples.

### RNA Seq

RNA sequencing was performed by Novogene (https://en.novogene.com).

### Gene function and pathway analyses

Pre-ranked gene set enrichment analysis (30, 31) was used to determine enriched pathways and ontology terms among high and low ATP genes. The gene list was collapsed to unique gene identifiers, and were ranked based on the magnitude of their ATP phenotype. The maximum gene set size was set at 500 genes, and the minimum size at 10 genes. Thousand random sample permutations were carried out using the Molecular Signature Database c2 v6.2 and c5 v6.2, and a significance threshold was set at a nominal *p*-value of 0.05.

### Principal component analysis

The R function prcomp was used for principal component analysis, with fractional labeling of metabolites as the input.

### Integrated metabolomics and transcriptomics analysis

Pathview (16, 32) was used to integrate metabolomic and transcriptomic data, with log2FC of genetic knockdowns versus controls as the input. For compound data input, we used log2FC of metabolite pool sizes versus controls averaged across all replicates. For gene input, we used log2FC of expression level versus controls averaged across all replicates.

## ACKNOWLEDGEMENTS

KN, NKB, were supported by the Joan and David Traitel Family Trust and NIH RO1NS091902 to KN. JLN and HN are supported by NIH/NCI award U54CA196519 and University of California Cancer Coordinating Committee Award to JN. HN was supported by a Japan Society for the Promotion of Science Fellowship.

## SUPPLEMENTARY DATA FILE LEGEND

**Data 1. *In vitro* and *in vivo* growth screen data, HCC827 cells, Source data for Figure 1**.

Mini-library CRISPRi screen in HCC827 cells, raw data. The readcounts for each sgRNA in all *in vitro* and *in vivo* (flank and orthotopic) replicates are included.

**Data 2. *In vitro* and *in vivo* growth screen data, H1975 cells, Source data for Figure 1**.

Mini-library CRISPRi screen in H1975 cells, raw data. The readcounts for each sgRNA in all *in vitro* and *in vivo* (flank and orthotopic) replicates are included.

**Data 3. Metabolomics C**^**13**^**-Glucose, Isotopologue Source data for Figure 3**.

**Data 4. Metabolomics C**^**13**^**-Glutamine, Isotopologue Source data for Figure 3**.

**Data 5. Metabolomics C**^**13**^**-Glucose, Fractional Contribution Source data for Figure 3**.

**Data 6. Metabolomics C**^**13**^**-Glutamine, Fractional Contribution Source data for Figure 3**.

**Data 7. RNAseq *in vitro* neg v *in vitro* sgc14orf2, Source data for Figure 2**.

**Data 8. RNAseq *in vitro* neg vs. *in vitro* sgMALSU1, Source data for Figure 2**.

**Data 9. RNAseq *in vitro* neg vs. *in vitro* sgTMEM261, Source data for Figure 2**.

**Data 10. RNAseq *in vivo* neg vs. *in vivo* sgMALSU1, Source data for Figure 2D**.

**Data 11. Pathway analysis of c14orf2, MALSU1, TMEM261: List of top 50 pathways**

## Supplementary Figures

**Supplementary Figure 1:**
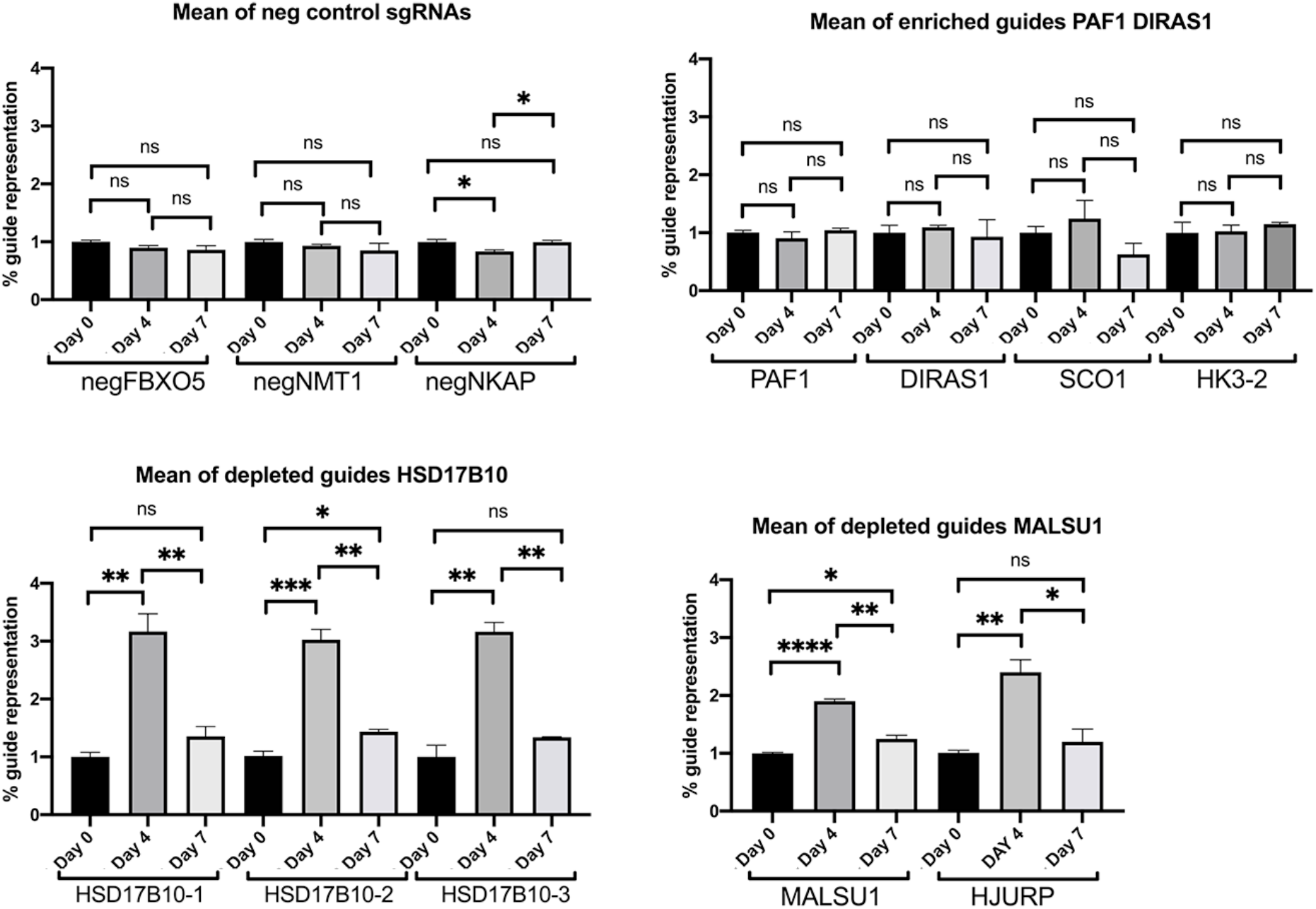
Initial sgRNA representation in HCC827 cells grown as flank tumors. HCC827 cells transduced with the mini-CRISPRi library were injected into the flanks of nude mice, then grown for either four or seven days (Day 4 or Day 7), after which tumors were removed and analyzed by sequencing and readcount analysis. Cells at injection are Day 0 samples. Number of replicates: Day 0 – n=3, Day 4 n=3, Day 7 n=2. sgRNA representation for selected sgRNAs (negative control sgRNAs, a subset of sgRNAs that were enriched in the initial flank experiment (day 28) (*PAF1, DIRAS1, SCO1, HK3*), and depleted sgRNAs in the flank experiment (*HSD17B10*, 3 individual sgRNAs, *MALSU1*, and *HJURP*) are plotted. Mean and s.e.m. shown for the labeled sgRNA at each timepoint (Day 0, Day 4 and Day 7). Student’s t-test, two sided performed for comparison of named sgRNA at each time point, * p<0.05, **p<0.01, ***p<0.001. Respiratory sgRNA representation are similar to non-respiratory control and glycolytic sgRNAs initially, enrich at day 4 or 7 relative to non-respiratory sgRNAs.

**Supplementary Figure 2.**
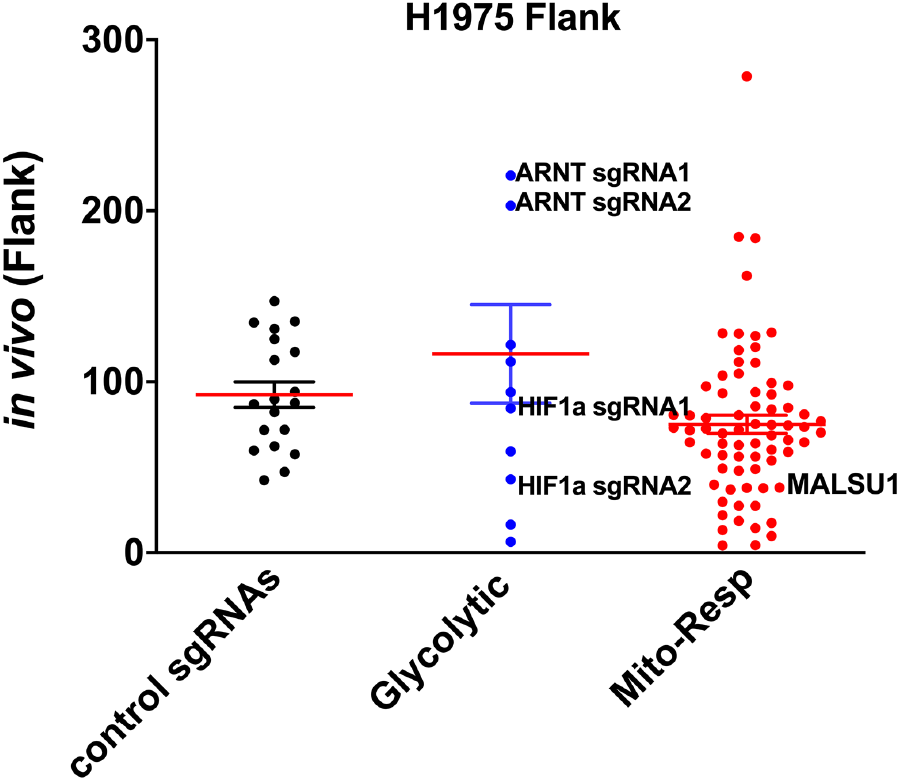
This figure relates to Figure 1. H1975 human *EGFR*-mutant lung cancer cells were transduced with the mini-CRISPRi library and then were injected into the subcutaneous space in flanks of nude mice and grown for 28 days. DNA from each tumor was sequenced and read counts for each sgRNA quantified. The read count for each sgRNA was normalized to negative control sgRNAs and the ratio of each sgRNA’s frequency in the tumor model relative to its frequency *in vitro* (immediately pre-injection). H1975 cells grown as flank tumors were analyzed n=6. Each dot represents a single sgRNA and indicates its average normalized representation displayed as control sgRNAs and sgRNAs targeting ATP-modulating genes classified as glycolytic or mito-respiratory (termed Mito-Resp). (Mean (red line) and s.e.m. shown, mean % representation of non-targeting, glycolytic and mito-respiratory sgRNAs are 92.5%, 116.3% and 75.0% respectively. One-way ANOVA of all three groups of sgRNAs demonstrate p value=0.03).

**Supplementary Figure 3.**
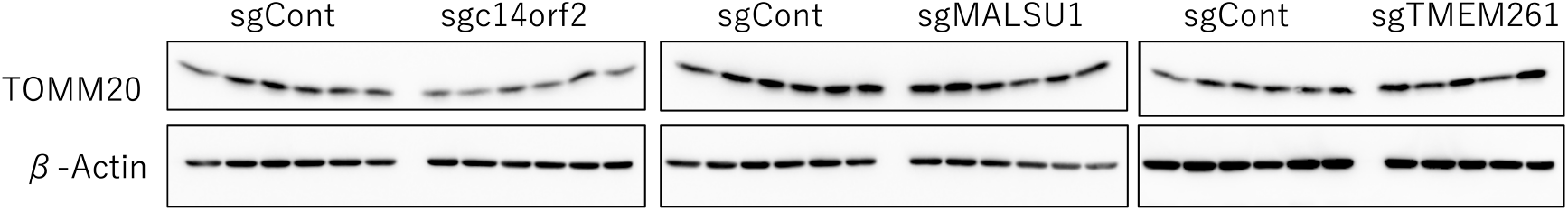
Mitochondrial protein level in HCC827 cells expressing mito-respiratory sgRNAs grown as flank tumors. Whole cell lysates were prepared from individual flank tumors (n=6 flank tumors of each sgRNA – control sgRNA (sgCont), *c14orf2, MALSU1, TMEM261*). Immunoblotting for the mitochondrial membrane protein TOMM20 was performed, with β-Actin as loading control.

**Supplementary Figure 4.**
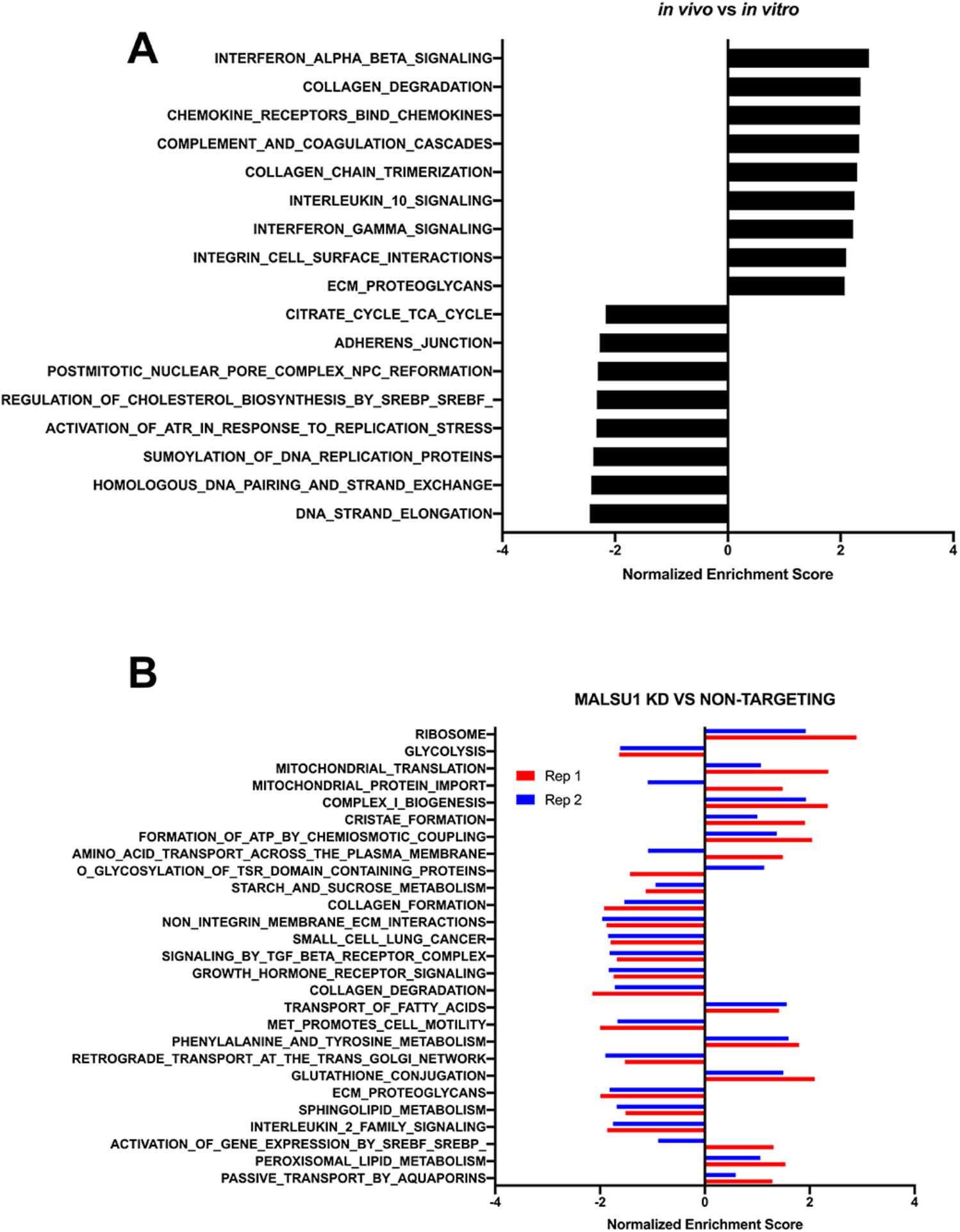
Pathway analysis of mito-respiratory silenced cells. HCC827 cells transduced with control sgRNA were injected into the flanks of nude mice, then grown for 28 days, after which tumors were removed and analyzed by RNA Seq (n=4 tumors analyzed). Gene set enrichment analysis was performed to compare control expression profiles in sgRNA-expressing cells grown *in vivo* to *in vitro* (A). B. The expression profile for each sgRNA *MALSU1* tumor replicate (Rep 1 and Rep 2, red and blue bars respectively) was analyzed, compared to control sgRNA and enriched/depleted pathways plotted (with normalized enrichment score). The replicates are concordant in pathway enrichment across most pathways, with the exception of mitochondrial protein import and amino acid transport across the plasma membrane.

**Supplementary Fig. 5.**
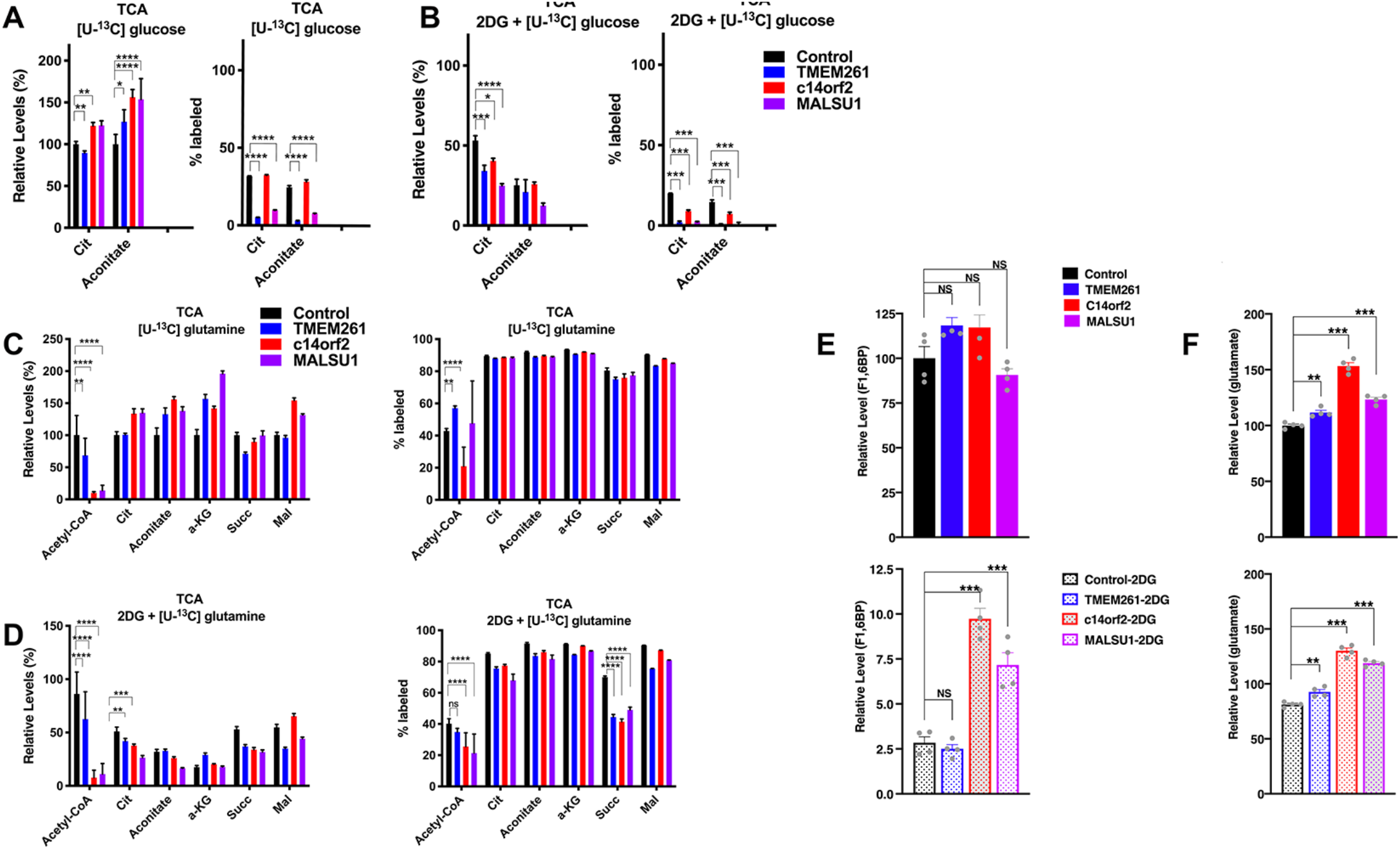
Relative levels and fractional labeling of glycolytic and TCA metabolites in mito-respiratory-silenced cells. HCC827 cells expressing individual CRISPRi sgRNA (control, or mito-respiratory hits *c14orf2, TMEM261, MALSU1*, silencing confirmation shown in Fig. 2A), were grown in basal media or media with 10 mM 2DG (n=4 per group) with either U-^13^C]glucose or [U-^13^C]glutamine for 18 hours. Cells were collected and metabolites analyzed. A. [U-^13^C]glucose labeled TCA intermediates (relative levels at left and % labeled at right) demonstrates that under basal conditions silencing of *TMEM261* and *MALSU1* is associated with significantly reduced citrate and aconitate (right panel, 1-way ANOVA *p<0.05, **p<0.01, ***p<0.001, ****p<0.0001). B. Forced respiration (2DG) reduces glucose-derived labeling of TCA intermediates, as expected. Mito-respiratory-silenced cells demonstrate significantly reduced citrate and aconitate compared to control cells. [U-^13^C]glutamine labeling of cells under basal (C) and respiratory (2DG)(D) conditions (Relative levels at left and % labeled at right). Under *basal* conditions, *TMEM261* demonstrates increased [U-^13^C]glutamine-derived labeling of acetyl-coA compared to control cells, whereas *c14orf2* cells demonstrate decreased [U-^13^C]glutamine-derived labeling of acetyl-coA compared to control cells. D. When 2DG is added, % labeled acetyl-coA is reduced all mito-respiratory cell lines (in the case of *TMEM261* cells, this reduction is relative to the elevated level under basal metabolism (C). In addition, all three mito-respiratory lines develop highly significant reductions in succinate labeling (right panel, *p<0.05, **p<0.01, ***p<0.001, ****p<0.0001). E/F. Relative levels of F1,6BP and glutamate are shown in cells grown without or with 2DG. These data relate to Figure 3A/B. One-way ANOVA with Dunnett’s multiple comparisons test *p<0.05, **p<0.01, ***p<0.001.

**Supplementary Figure 6.**
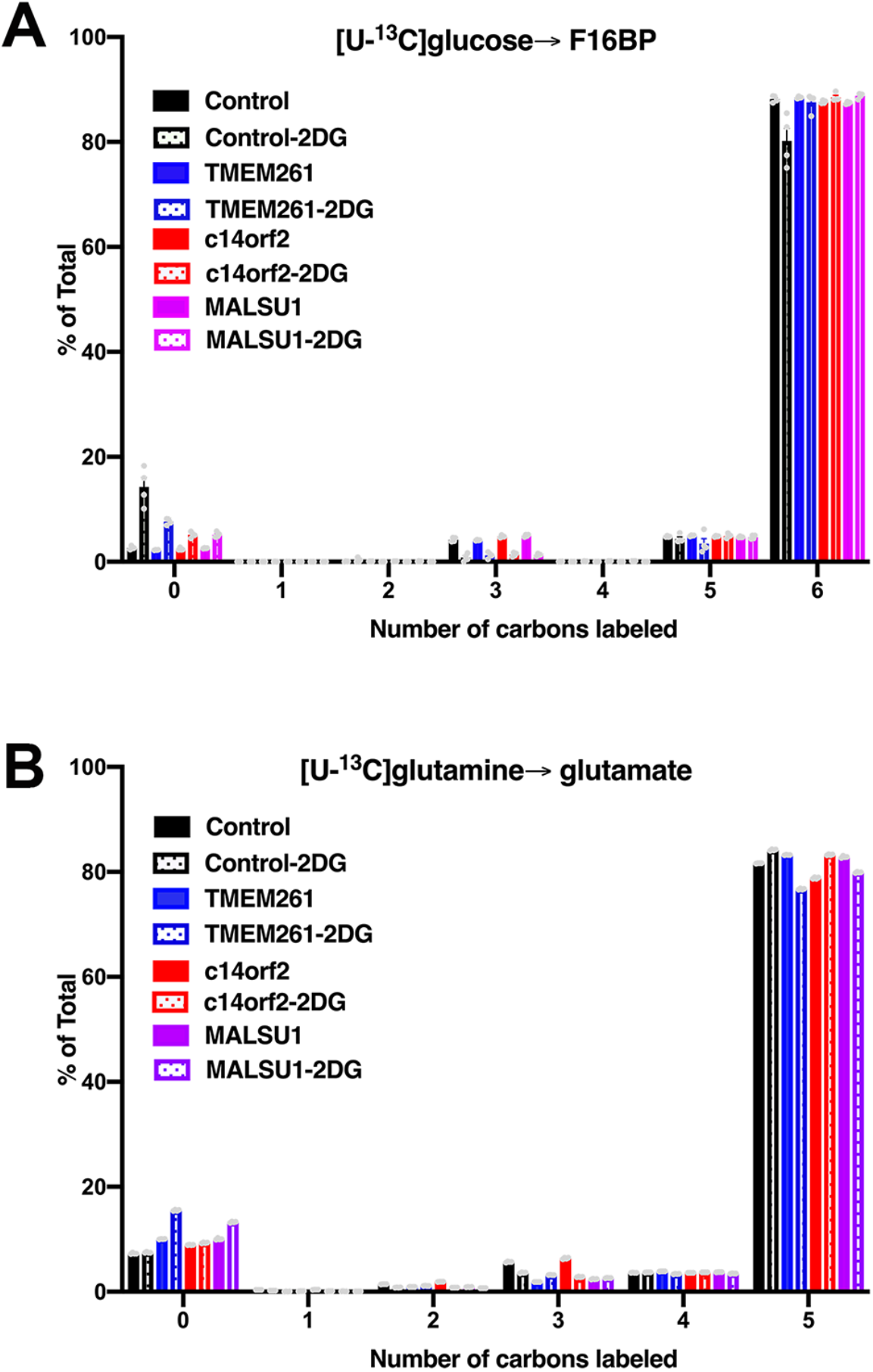
Isotopologues for F16BP and Glutamate. A. All labeled isotopologues for ^13^C glucose→ F16BP indicates % total unlabeled (0 carbon) or completely labeled (6 carbon) (n=4 replicates per sample, mean and sem shown). This data relates to Figure 3A. B. All labeled isotopologues for ^13^C glutamine→ Glutamate indicates % total unlabeled (0 carbon) or completely labeled (5 carbon) (n=4 replicates per sample, mean and sem shown). This data relates to Figure 3A.

**Supplementary Figure 7.**
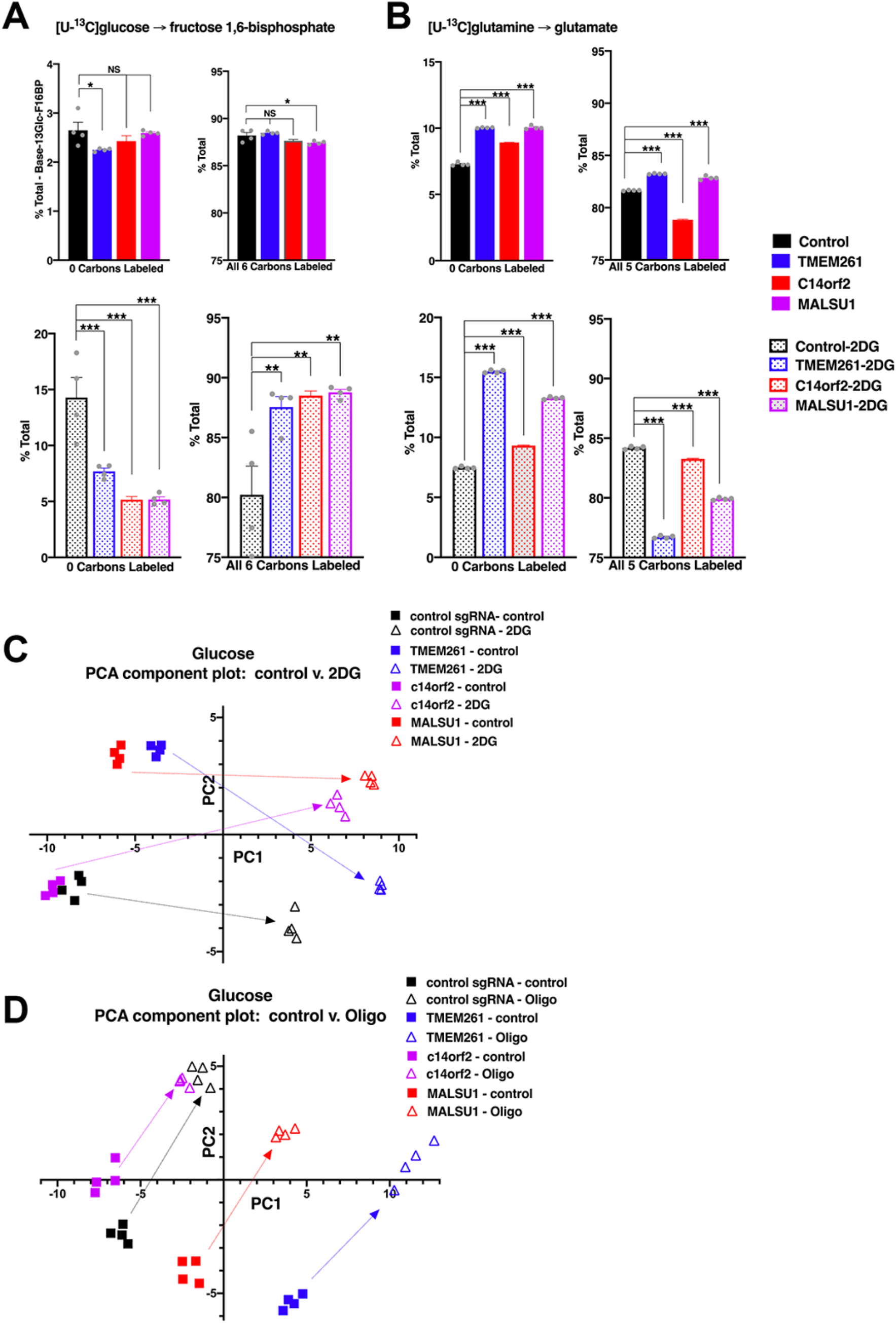
Metabolomics analysis and Principal Component Analysis. HCC827 cells expressing individual CRISPRi sgRNA (control, or mito-respiratory hits *TMEM261, c14orf2, MALSU1*, silencing confirmation shown in Fig. 2A), were grown in basal media or media with 10 mM 2DG (n=4 per group, individual data points shown as grey dots) with either [U-^13^C]glucose or [U-^13^C]glutamine for 18 hours. Cells were collected, metabolites extracted and analyzed by mass spectrometry. **A**. [U-^13^C] glucose→ fructose 1,6-bisphosphate indicates % total unlabeled (0 carbon) or completely labeled (6 carbon) with top graphs indicating percent total and fractional labeling (left and right, respectively) under basal conditions and the bottom graphs indicating percent total and fractional labeling in the presence of 2DG (n = 4, mean, sem). Respiratory-deficient cells demonstrate relative resistance to 2DG, maintaining glycolytic activity and labeling in the presence of 2DG (shown in bottom right panel, 1-way ANOVA, Dunnett’s multiple comparisons test, *p<0.05, **p<0.01, ***p<0.001, NS-not significant). (n=4 replicates per sample, mean and sem shown) **B**. [U-^13^C]glutamine→ glutamate analysis indicates total unlabeled (0 carbon labeled) on the left and fully labeled (all 5 carbons labeled) on the right. Control cells maintain glutamine uptake (left panel) after addition of 2DG, which is not expected to affect respiration. Compared to control, respiratory-deficient cells demonstrate reduced uptake of glutamine (reflected in increased total unlabeled) at baseline (left panel). Adding 2DG exaggerates this defect in glutamine uptake by respiratory-deficient cells (reflected in increased % unlabeled, shown in left panel). In the right panel, U-^13^CGlutamine→ glutamate indicates activity through TCA. 2DG increases glutamine metabolism through TCA in control cells, consistent with compensatory metabolic shift to TCA and intact TCA. Basally, respiratory-deficient cells (*TMEM261* and *MALSU1*) demonstrate a modest but significant increase in TCA metabolism compared to control sgRNA cells (right panel), which we hypothesize represents a compensatory drive to increase activity through TCA to accommodate the respiratory defect. However, 2DG overcomes this compensation and further exacerbates the TCA defect in respiratory-deficient cells, which demonstrate significantly and severely decreased metabolism through TCA. **C/D**. Principal component analysis was applied to the fractional contribution values of the metabolomics data for HCC827 cells expressing individual CRISPRi sgRNA (control, or mito-respiratory hits *TMEM261, MALSU1*, or *c14orf2*. ^13^C glucose-derived labeling of cells grown under either control or 2DG media was compared in **C**, control or oligomycin in **D**. The PC1 and PC2 values of the fractional contribution analysis is plotted for each cell line grown under either control or 2DG conditions (n=4 replicates for each line) is plotted.

**Supplementary Figure 8.**
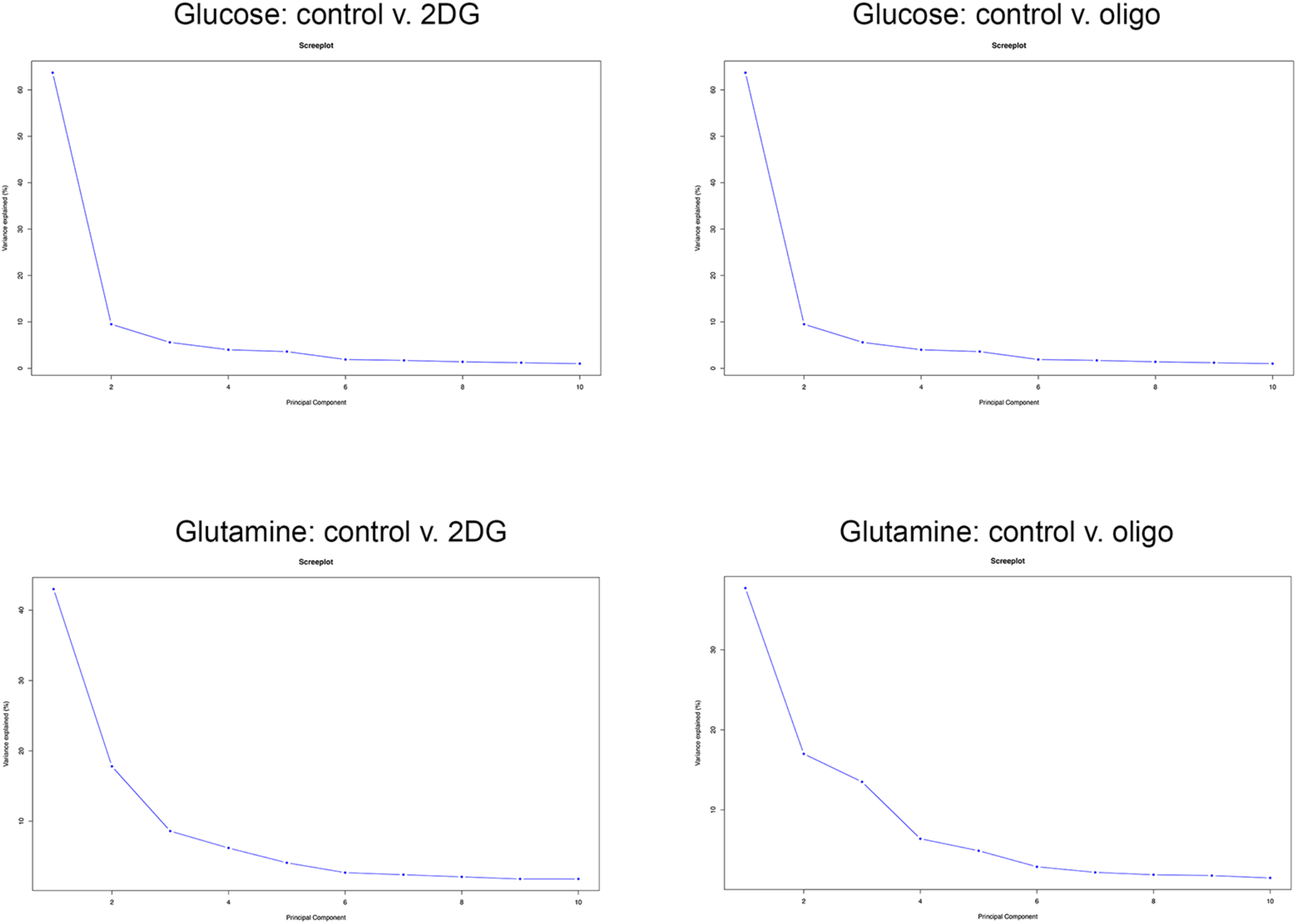
Skreeplots of principal component analysis of ^13^C-glucose or ^13^C-glutamine metabolomics. Principal component analysis (PCA) was performed to assess similarities in metabolite responses among the mito-respiratory cell lines. All four cell lines (control sgRNA and three mito-respiratory lines) were pooled to analyze either ^13^C-glucose or ^13^C-glutamine metabolites under control conditions and either 2DG or oligomycin. Two principal components explain the majority of the variance for all four comparisons.

**Supplementary Figure 9.**
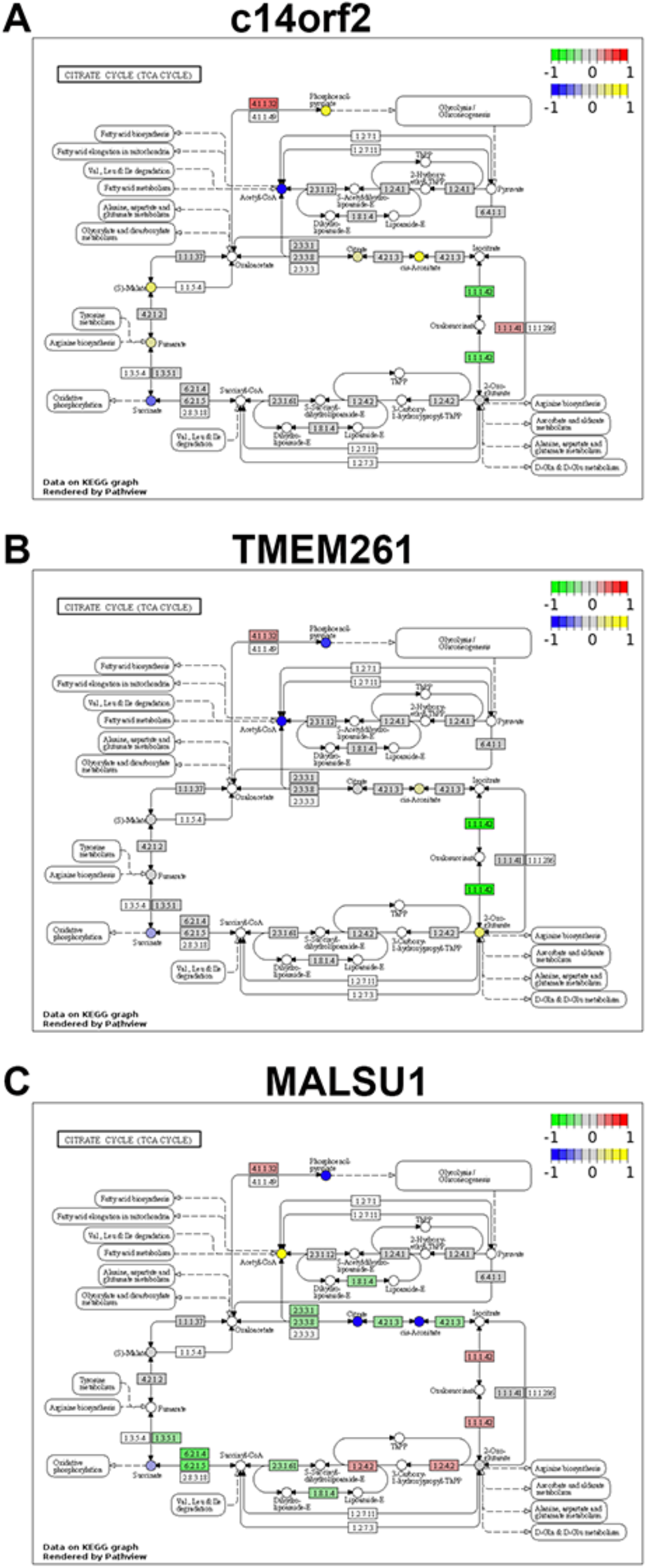
Integrated metabolomics and transcriptomics-based pathway analysis distinguishes metabolic network structure among mito-respiratory cells. Pathway analysis was performed by integrating transcriptome and metabolomics data for HCC827 cells expressing individual CRISPRi sgRNA (control or mito-respiratory hits TMEM261, MALSU1, or c14orf2) grown *in vitro* and plotted on a KEGG graph depicting pathway components for citrate cycle (TCA cycle).

